# DNA binding by ATPase-adjacent domains stimulates SMCHD1 ATPase activity

**DOI:** 10.64898/2026.02.05.704097

**Authors:** Alexandra D. Gurzau, Richard W. Birkinshaw, Andrew Leis, Tamara Cameron, Megan Iminitoff, Kelsey Breslin, Iromi Wickramasinghe, Samantha Eccles, Hannah K. Vanyai, Quentin A. Gouil, Peter E. Czabotar, James M. Murphy, Marnie E. Blewitt

## Abstract

SMCHD1 is an epigenetic regulator in which heterozygous variants are reported in facioscapulohumeral muscular dystrophy (FSHD), as well as Bosma arhinia microphthalmia syndrome (BAMS). While we have previously shown that SMCHD1 is able to interact with nucleic acids via its hinge domain, we have now identified a second DNA-binding site that is located C-terminal to the ATPase domain and formed by two domains: the Bromo-adjacent homology (BAH) and immunoglobulin-like 1 (IGL-1) domains. Here, we report their mode of DNA-interaction and we present the first high-resolution structure of the wild-type human SMCHD1 ATPase using site-directed mutagenesis and structural analysis via cryo-EM. We also reveal that DNA-binding at the BAH-IGL-1 domains stimulates the ATPase activity of full-length SMCHD1 *in vitro*, and demonstrate in a mouse model that ATP hydrolysis is essential for SMCHD1 function *in vivo*. Together, these findings establish bidentate DNA binding and DNA-stimulated ATP hydrolysis as central features of SMCHD1 function, providing new mechanistic insight into how SMCHD1 regulates gene silencing.

**GRAPHICAL ABSTRACT:** 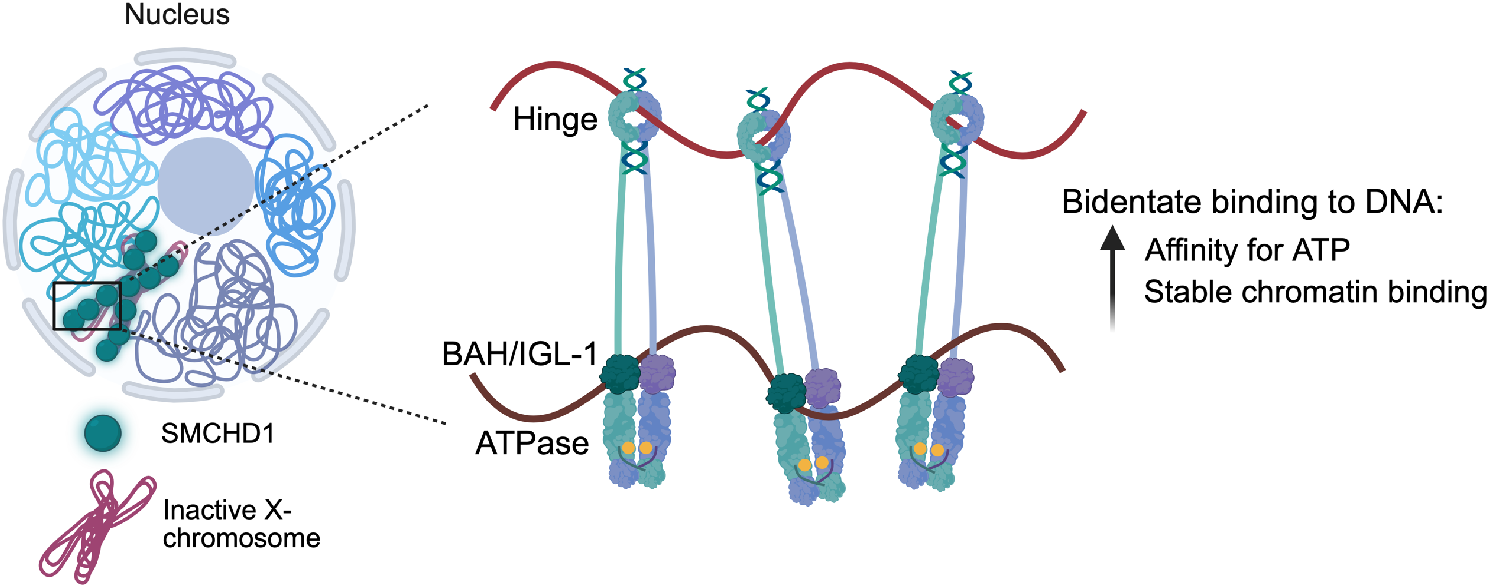

## INTRODUCTION

Structural Maintenance of Chromosomes Hinge Domain-containing 1 (SMCHD1) is an epigenetic regulator that plays a pivotal role in X-chromosome inactivation as well as in the regulation of various autosomal genes, including selected imprinted genes (1,2). Variants in the human *SMCHD1* gene are implicated in two debilitating conditions: facioscapulohumeral muscular dystrophy (FSHD) (3) and the rare craniofacial disorder, Bosma arhinia microphthalmia syndrome (BAMS) (4). While we previously showed that FSHD-and BAMS-associated mutations in *SMCHD1* alter the protein’s catalytic rate (5), the molecular mechanisms by which SMCHD1 mediates gene silencing and how pathogenic variants disrupt this process are not fully understood. There is therefore a growing interest in determining how SMCHD1 functions both in normal development and in disease.

The *SMCHD1* gene encodes a ∼230-kDa nuclear protein that is conserved across vertebrates, and consists of an N-terminal GHKL-type ATPase (6-8) and a C-terminal SMC hinge domain (9-11), connected by a long intermediate region of unknown structure or function. Over the last few years, studies on these two isolated domains have revealed more detailed structural and functional information about SMCHD1 (7-11). However, the high-resolution structure of the full-length protein, its mechanism of action, as well as its mode of recruitment to chromatin are all largely unknown. In particular, SMCHD1’s ATPase activity has emerged as a key regulatory element, whose functional role and governing mechanisms remain to be determined. We previously demonstrated that loss-of-function FSHD-associated mutations in *SMCHD1* that are located both within the ATPase region and immediately downstream frequently lead to reduced catalytic activity (5), indicating that ATPase activity may be regulated by other regions of the protein, and that ATP turnover is a crucial aspect of SMCHD1 function.

The functional importance of ATP hydrolysis in SMCHD1 has been explored recently through studies of the E147A point mutation, which targets a catalytic residue that is conserved across members of the GHKL family of proteins and is necessary for the ATP hydrolysis step (7,8). At the protein level, this mutation promotes constitutive dimerization of the ATPase domains upon ATP binding (8). At the cellular level, the E147A mutation disrupts SMCHD1 accumulation on the inactive X chromosome (Xi) and compromises its silencing (6,12). Live-cell imaging studies suggest that while ATP hydrolysis is not strictly required for chromatin engagement, it is necessary for stable, long-term SMCHD1 interactions, particularly on the Xi (12). Nevertheless, how ATPase activity is mechanistically coupled to chromatin association and gene repression remains unclear.

In this study, we investigated the role of SMCHD1 ATPase activity in gene regulation, and explored how regions outside the ATPase domain contribute to SMCHD1 function. We generated a novel mouse model and found that the E147A mutation induces a *Smchd1*-null effect *in vivo*, indicating that ATPase function is essential for SMCHD1’s role during early development. Using biochemical and structural approaches, we defined a novel DNA-binding site within the SMCHD1 protein formed by the BAH and IGL-1 domains, and determined their structures using cryo-electron microscopy (cryo-EM). We additionally demonstrated that SMCHD1’s ATPase activity is stimulated by DNA interacting with the BAH-IGL-1 domains, and that this second DNA binding site is required for normal chromatin localisation. Together these data provide new insight into the molecular mechanisms underlying SMCHD1-mediated gene repression.

## MATERIALS AND METHODS

### Mouse strains and genotyping

Mice were bred and maintained under standard animal husbandry conditions. All protocols were approved by the WEHI Animal Ethics Committee, under animal ethics approval numbers 2018.004, 2020.050, 2020.048 and 2023.050 and 2023.045. Mice were kept in individually ventilated cages on standard chow, in a 14h/10h light/dark cycle. *Smchd1*^GFP/GFP^ mice were maintained on an inbred C57BL/6 background as previously described (13).

The Smchd1 E147A mutation was introduced onto the *Smchd1*^GFP/GFP^ background by microinjection of two guide RNAs and a homology repair vector (sequences in **Supplementary Table 1**) along with Cas9 protein, into single cell fertilised eggs (14). The two guide RNAs enabled an exon replacement approach to introduce the point mutation. The repair vector also introduced silent point mutations at the PAM sites to prevent more than one cut with Cas9. This work was conducted by the MAGEC facility at WEHI. The new allele was backcrossed to C57BL/6 (with or without the *Smchd1*^GFP^ allele) for more than five generations before experiments began. Therefore, the *Smchd1*^GFP-E147A^ allele was produced and maintained on a C57BL/6 strain background. The *Xist*^ΔA^ allele (15) was maintained on a 129P4 background (16).

The Castaneus mice were purchased from Jackson labs and maintained as a closed colony. The E147A allele was genotyped initially by PCR and sequencing, then once established using allelic discrimination assays (primers and probes in **Supplementary Table 1**). The *Smchd1*^GFP^ allele is genotyped by PCR (13) (primers in **Supplementary Table 1**). The sex of embryos was determined using PCR, with primers that detect the X-linked gene *Otc* and the Y-linked gene *Zfy* (primers in **Supplementary Table 1**). The *Xist*^ΔA^ allele is genotyped using PCR (16) (primers in Supplementary Table 1).

### Generation of Neural Stem Cells

Neural stem cells were generated as we have previously described (11,13,16). Briefly, we set up timed matings, and dissected embryos at embryonic day 14.5. The cortices of each embryo were dissociated and grown as an adherent monolayer on laminin coated tissue culture treated plates. Their growth media was NeuroCult NSC Basal Medium (mouse) with NeuroCult Proliferation Supplement (Stem Cell Technologies), with the addition of recombinant human EGF (20 ng/mL) and recombinant human basic FGF (20ng/mL, both from Stem Cell Technologies).

### Neural stem cell samples for RNA sequencing

To enable us to assess the X inactivation status in neural stem cell populations, we created neural stem cells from embryos that were *Smchd1*^GFP-E147A/+^; *Xist*^ΔA/+^ or *Smchd1*^+/+^; *Xist*^ΔA/+^ for females, or *Smchd1*^GFP-E147A/+^; *XY* or *Smchd1*^+/+^; *XY* for males. In addition to the genetic changes, the animals were all F1, derived from a Castaneus father mated with a mother that was 129/C57 F1. Therefore, all Castaneus alleles were of paternal origin, and for the X inactivation analyses the *Xist*^ΔA^ allele ensured that the Castaneus X was always the inactive X chromosome. This approach allowed allele-specific analysis of the RNA sequencing data.

### RNA sequencing analysis and data accessibility

Libraries were prepared using the Illumina TruSeq V2 stranded mRNA sample preparation kits from 500 ng total RNA as per manufacturer’s instructions. Fragments above 200 bp were size-selected and cleaned up using AMPure XP magnetic beads. Final cDNA libraries were quantified using D1000 tape on the TApeStation (4200, Agilent Technologies) for sequencing on the Illumina Nextseq500 platform using 80 bp, paired-end reads.

RNA-seq reads were trimmed for adapter and low-quality sequences using TrimGalore! V0.4.4, before mapping onto the GRCm38 mouse genome reference with HISAT2 v2.0.5 (17) in paired-end mode and disabling soft-clipping. Because the samples have 129/SvJ maternal alleles and either Cast/EiJ or C57BL/6 paternal alleles (except the paternal X that is only Cast), we prepared N-masked versions of the genome using SNPsplit v0.3.2. For autosomal gene expression analysis, both 129/SvJ and Cast/EiJ SNPs are N-masked. For X-linked expression analysis, we used a dual-hybrid genome prepared with SNPsplit. Alignments specific to the C57BL/6, 129/SvJ and Cast/EiJ alleles were separated using SNPsplit v0.3.2 in paired-end mode.

Gene counts were obtained in R 3.5.1 (R development Core Team 2019) from the split and non-split bam files with the featureCounts function from the Rsubread package (1.32.1) (18,19), provided with the GRCm38.90 GTF annotation downloaded from Ensembl, and ignoring multi-mapping or multi-overlapping reads.

For analysis of global expression changes, total (non-haplotyped) counts were normalized in edgeR v3.24.0 (20,21) with the TMM method (22). Genes were defined as expressed and retained for differential expression analysis if they had at least one count per million (cpm) in at least one third of the libraries. Differential expression analysis was performed with likelihood ratio tests. P-values were corrected with the Benjamini-Hochberg method (23). Differential expression results were visualized with Glimma 1.10.0 (24), with differential expression cut-offs of adjusted p-value<0.05. Plots were generated in R with the ggplot2 (25), cowplot (26), ggbeedswarm (27), and ggrepel (28) packages.

For allele-specific analysis of autosomal imprinted genes, we genotyped samples as previously (29), fitting a recursive partition tree with the rpart function (R Development Core Team, 2019; (30)) to the proportion of Cast/EiJ or C57BL/6 reads in each 100 kb bin tiling the genome, with options minsplit = 4, cp = 0.05. We compiled a list of all 316 known mouse imprinted genes and kept genes with at least 20 haplotyped counts in at least 2 samples per genotype. To determine whether these genes were imprinted in the neural stem cells, we then fitted a logistic regression with the glm function from the stats package (R Development Core Team, 2019) on the maternal and paternal counts for each gene in the wild-type samples and retained genes with an absolute log-ratio of expression greater than or equal to log(1.5). A betabinomial test (from the aod package, (31)) was performed for each gene to assess for differences in parental expression ratios between wild-types and mutants. The p-values were corrected with the Benjamini-Hochberg method.

To investigate the effect of the E147A Smchd1 mutation on X-chromosome inactivation in females, we used edgeR’s paired-sample design (32) with common dispersion to model the maternal (Xa) and paternal (Xi) counts as a function of genotype. We tested the effect of genotype with a likelihood ratio test. The p-values were corrected with the Benjamini-Hochberg method. We used the criteria of 5% false discovery rate (FDR).

### Protein expression

The DNA sequence encoding human SMCHD1 residues 571-688, 688-819 and 571-819 were cloned into a pPROEX HTb vector (Invitrogen) and proteins were expressed in *E. coli* BL21-Codon Plus (DE3) cells (Agilent). Mutations were introduced by oligonucleotide-directed mutagenesis PCR (**Supplementary Table 1**) and insert sequences verified by Sanger sequencing (AGRF). Briefly, cells were cultured in Super broth at 37°C under shaking conditions at 200 rpm, until an OD_600_ ∼0.6-0.8 was reached. The temperature was then reduced to 18°C and protein expression was induced with the addition of 0.5 mM IPTG overnight. Human SMCHD1 constructs encompassing residues 25-819 and 25-1959 (and related point mutations) were cloned into a pFastBac Htb (Invitrogen) vector, for expression via the Bac-to-Bac (Invitrogen) insect cell expression system, which consisted of generating recombinant bacmids in *E. coli* DH10MultiBac (ATG Biosynthetics) cells using established protocols (33). Both the bacterial and insect cell expression constructs encode an N-terminal TEV protease-cleavable 6x Histidine tag for Ni^2+^ affinity purification. For insect cell expression, bacmid transfection and P2 virus generation was performed using *Sf*21 cells suspended in Insect-XPRESS protein-free medium with L-glutamine (Lonza) following the FuGENE HD Transfection Reagent Protocol (Promega). On day 0, 12 mL P2 virus was used to transduce Expi*Sf*9 cells suspended in 1 L Expi*Sf*9 CD Medium (Thermo Fisher Scientific) at 6 x 10^6^ cells/mL. The culture was incubated in a 3L Fernbach flask with vented cap (Corning) at 27.5°C, shaking at 100 rpm. On day 4, cells were harvested by centrifugation at 500 x*g* for 5 min at room temperature, and snap-frozen in liquid nitrogen for storage at -80°C.

### Protein purification

*Sf*21 insect cell or bacterial cell pellets were resuspended in lysis buffer (0.5 M NaCl, 20 mM Tris–HCl pH 8.0, 20% (v/v) glycerol, 5mM imidazole pH 8.0, 0.5mM TCEP), supplemented with 1mM PMSF and 1X cOmplete EDTA-free protease inhibitor (Roche). Cells were disrupted by sonication while maintaining the lysate at 4°C, and insoluble material was removed by centrifugation at 45,000 xg for 30 min at 4°C. Supernatants were subjected to Ni-chromatography (cOmplete His-Tag purification resin, Roche) and following extensive washing, eluted in lysis buffer containing 250 mM imidazole pH 8.0. Following overnight cleavage of the His-tag with TEV protease, proteins were passed back over fresh Ni^2+^ resin and the unbound fraction was collected. These were further concentrated and purified by either Superose 6 increase 10/300 GL (Cytiva) size-exclusion chromatography for the 25-1959 SMCHD1 protein (150 mM NaCl, 10 mM HEPES pH 8.0, 0.5 mM TCEP and 10% glycerol) or for all remaining SMCHD1 constructs, subjected to size-exclusion chromatography on a Superdex-200 10/300 GL size exclusion chromatography (Cytiva) with elution in 100 mM NaCl, 20 mM HEPES pH 7.5, 0.5 mM TCEP. Protein purity was evaluated by reducing SDS– PAGE with Stain-Free visualization (Biorad) and fractions of interest were pooled, snap-frozen in liquid nitrogen and stored at −80°C until required.

### Mass Photometry

Data acquisition was performed on a Refeyn Two^MP^ mass photometer (Refeyn Ltd) using Acquire MP (Refeyn Ltd), using protein concentrations ranging between 20-100 nM. Calibration with protein samples of known molecular weight was initially performed. Each movie was recorded for 60 seconds and processed and analysed using DiscoverMP (Refeyn Ltd).

### Cryo-electron microscopy (cryo-EM) sample preparation

Purified wild-type human SMCHD1 protein encompassing residues 25-819, or purified 25-688 residue E147A human SMCHD1 protein, were incubated with 2-fold molar excess of AMPPNP and MgCl_2_ for 2 hours on ice, followed by a further size-exclusion chromatography step on a Superdex 200 increase 10/300 GL (Cytiva) column to separate monomeric and dimeric populations. The dimer peak was collected and vitrified by applying 4 μL at 0.75 mg/mL to UltrAuFoil R1.2/1.3 grids (Quantifoil), which were glow discharged for 180 seconds at 25 mA in a GloQube apparatus (Quorum). Vitrification was performed using a Vitrobot Mark IV (Thermo Fisher Scientific) at 4°C and 100% humidity with a nominal blot force zero for 4 seconds. The resulting grids were imaged on a Titan Krios G4 microscope operated at 300 keV equipped with a Falcon 4 detector (Thermo Fisher Scientific), at x96,000 nominal magnification, corresponding to a pixel size of 0.808 Å at the detector using EPU automated acquisition software Version 3.11 (Thermo Fisher Scientific).

### Cryo-EM image processing and model building

Image processing was performed using cryoSPARC (34). The movies were aligned and averaged using patch motion correction, and contrast transfer function parameters were estimated using Patch CTF estimation. Particles were picked using Blob Picker, and the particle coordinates were used for 2D class averaging. Selected 2D classes were then used to create an *ab initio* model (**Fig. S3, S4)**.

### Fluorescence polarization DNA-binding assays

25 nM of 6’-Fam labelled single- or double-stranded DNA (Integrated DNA Technologies), with sequences outlined in **Supplementary Table 1**, were incubated with recombinant SMCHD1 protein at final concentrations ranging between 0-100 µM. Protein samples were diluted in protein sample buffer (SEC buffer) in 10 µL reactions, as described previously (10). Reactions were set up in 384-well low flange black flat bottom plates (Corning) in duplicate and incubated at room temperature for 20 minutes in the dark. Emission polarization values were measures with an Envision plate reader (PerkinElmer) using a 480 nm excitation filter, a 535 nm static and polarized filter and FITC FP dual mirror. Results were plotted using Prism 10.

### Electromobility shift assays (EMSA)

EMSA was performed as using the protocol of Griese *et al*. (35). Briefly, 50 nM of 6-Fam-labelled oligonucleotides (Integrated DNA Technologies) as listed in **Supplementary Table 1**, were mixed with recombinant SMCHD1 protein in varying molar excess amounts over DNA in EMSA buffer (75 mM NaCl, 20 mM Tris pH 7.5, 5% glycerol, 0.005% Triton X-100, 0.5 mM MgCl_2_) in a total volume of 10 μL. Samples were incubated for 30 minutes at room temperature, following the addition of 2.5 μL of 50% (v/v) glycerol. The samples were loaded either onto a DNA retardation gel (6%) 1.0 mM (Thermo Fisher Scientific) or onto a homemade 0.5% agarose gel in 0.5x TBE buffer [890 mM Tris base, 890 mM boric acid, 20 mM EDTA (pH 8.0)] and separated for 90 minutes at 90 V, after which they were either stained with SYBRgold (1:20,000 in milliQ) for 30 min at room temperature and destained with milliQ for 15 minutes and scanned on a ChemiDoc imaging system (BioRad), or visualised using the 6’FAM fluorophore on the DNA probe using an Amersham Typhoon Biomolecular Imager.

### Fluorescence polarization ATPase assays

10 µL reactions were set up in triplicated in 384-well low flange, black, flat-bottom plates (Corning) containing 7 µL reaction buffer (50 mM HEPES pH 7.5, 4 mM MgCl_2_, 2 mM EGTA), 1 µL recombinant protein at various concentrations or SEC buffer control, 1 µL nuclease-free water or DNA (ssDNA or dsDNA at a final concentration of 0.5 µM) and ranging concentrations of ATP substrate, as outlined previously (5). Reactions were incubated at room temperature for 1 hour in the dark and stopped by the addition of 10 µL Stop&Detect buffer (1X Detection buffer, 4nM ADP Alexa Fluor 633 tracer, 128 µg/mL ADP^2^ antibody) and incubated for a further 1 h in the dark, at room temperature. Fluorescence polarization readings (mP) were measured on an Envision plate reader (PerkinElmer Life Sciences) fitted with excitation filter 620/40 nm, emission filters 688/45 nm (s and p channels) and a D658/fp688 dual mirror. Concentrations of ADP produced by the reactions were estimated by a 12-point standard curve, as outlined in the manufacturer’s protocol (Transcreener ADP^2^ FP assay, BellBrook Labs), and data were plotted and analysed in Prism 10.

### Surface Plasmon Resonance (SPR)

Experiments were performed in HBS-EP buffer consisting of 10 mM HEPES pH 7.4, 150 mM sodium chloride, 3.4 mM EDTA, 0.05% Tween-20 and 1 mM TCEP, at 25 ºC, on a BIAcore S200 using a SA sensor immobilized with biotinylated 60 base-pair single- or -double stranded DNA probes synthesised by IDT (sequences in **Supplementary Table 1**). Binding affinities were determined by flowing purified SMCHD1 protein (residues 571-819) in a concentration range of 0.625-20 µM. Samples were injected in a multi-cycle run (flow rate 30 µL/min, contact time 120 s, dissociation 600 s). Steady state binding data were fitted to a 1:1 binding model and evaluated in a BIAevaluation software. Data were plotted using Prism 10.

### HEK293T cell culture

HEK293T cells were cultured in DMEM (Gibco: 11965092) containing 10% (v/v) fetal bovine serum (FBS) and 100 U/ml penicillin and 100 µg/mL streptomycin (Gibco: 15140122). Cells were grown at 37°C in 5% CO_2_.

### CRISPR/Cas9-mediated mutagenesis of SMCHD1 in HEK293T cells

To insert single amino acid mutations in SMCHD1 in HEK293T cells, we designed guide RNA and homology-directed repair (HDR) donor templates (**Supplementary Table 1**) using the IDT Alt-R CRISPR-Cas9 guide RNA and HDR Design tools. Genome editing was performed using a ribonucleoprotein CRISPR/Cas9 editing approach, where reagents were electroporated into wild-type HEK293T cells using a nucleofection kit (SF Cell Line 4D-Nucleofector X Kit S; Lonza: V4XC-2032). We included a ‘no guide RNA’ control for each nucleofection reaction, but which consists of the Cas9 protein and HDR template. Multiple clonal cell lines were then established for each mutation by limited dilution. Editing efficiency was determined on these clonal cell line populations by MiSeq DNA sequencing (Illumina) after PCR amplification of the Cas9-targeted region and secondary PCR using overhang sequences (**Supplementary Table 1**).

### Immunofluorescence microscopy

Immunofluorescence microscopy studies were performed using an established protocol (36). Cells were washed in phosphate-buffered saline (PBS) and fixed in PBS, 3% (w/v) paraformaldehyde for 10 min at room temperature. Cells were washed three times with PBS for 5 min each and then permeabilized on ice with 0.5% (v/v) Triton X-100 in PBS, followed by three washes in PBS for 5 min each. Cells were blocked in 1% (w/v) bovine serum albumin (BSA; Life Technologies) for 15 min, followed by a 45-min incubation in a dark and humid chamber at room temperature with a primary anti-Smchd1 antibody (in-house monoclonal clone 2B8) and anti-H3K27me3 Alexa-Fluor-647-conjugate antibody (Cell Signalling Technology:C36B11), diluted 1:100 in 1% (w/v) BSA. Cells were washed three times in PBS for 5 min each and incubated for 40 min at room temperature in a dark and humid chamber with a secondary anti-rat 568 antibody (Life Technologies: A-11077) diluted 1:500 in 1% (w/v) BSA. Cells were washed three times in PBS for 5 min each and stained with 4′,6-diamidino-2-phenylindole (DAPI) for 10 min at room temperature, followed by another two washes with PBS. Coverslips were mounted in VECTASHIELD H1000 mounting medium (Vector Laboratories). Cells were visualized with a Zeiss LSM 880 NLO microscope at ×63 magnification, and *z* stacks were acquired. Images were analysed with the open-source ImageJ distribution package (FIJI). Statistical analyses were calculated and plotted with Prism 10, using an unpaired t-test with Welch’s correction.

## RESULTS

### The E147A allele phenocopies *Smchd1*-deficiency in a knockin mouse model

To further understand the catalytic-dependent functions of SMCHD1, we generated a new mouse line carrying the E147A mutation that we and others have previously shown reduces the ATP hydrolysis activity of SMCHD1 (7,8). We introduced the mutation into a *Smchd1*^GFP^ allele (13) to create *Smchd1*^GFP-E147A^ and allow tracking of SMCHD1 using the GFP tag (**Fig. 1A**).

**Figure 1.**
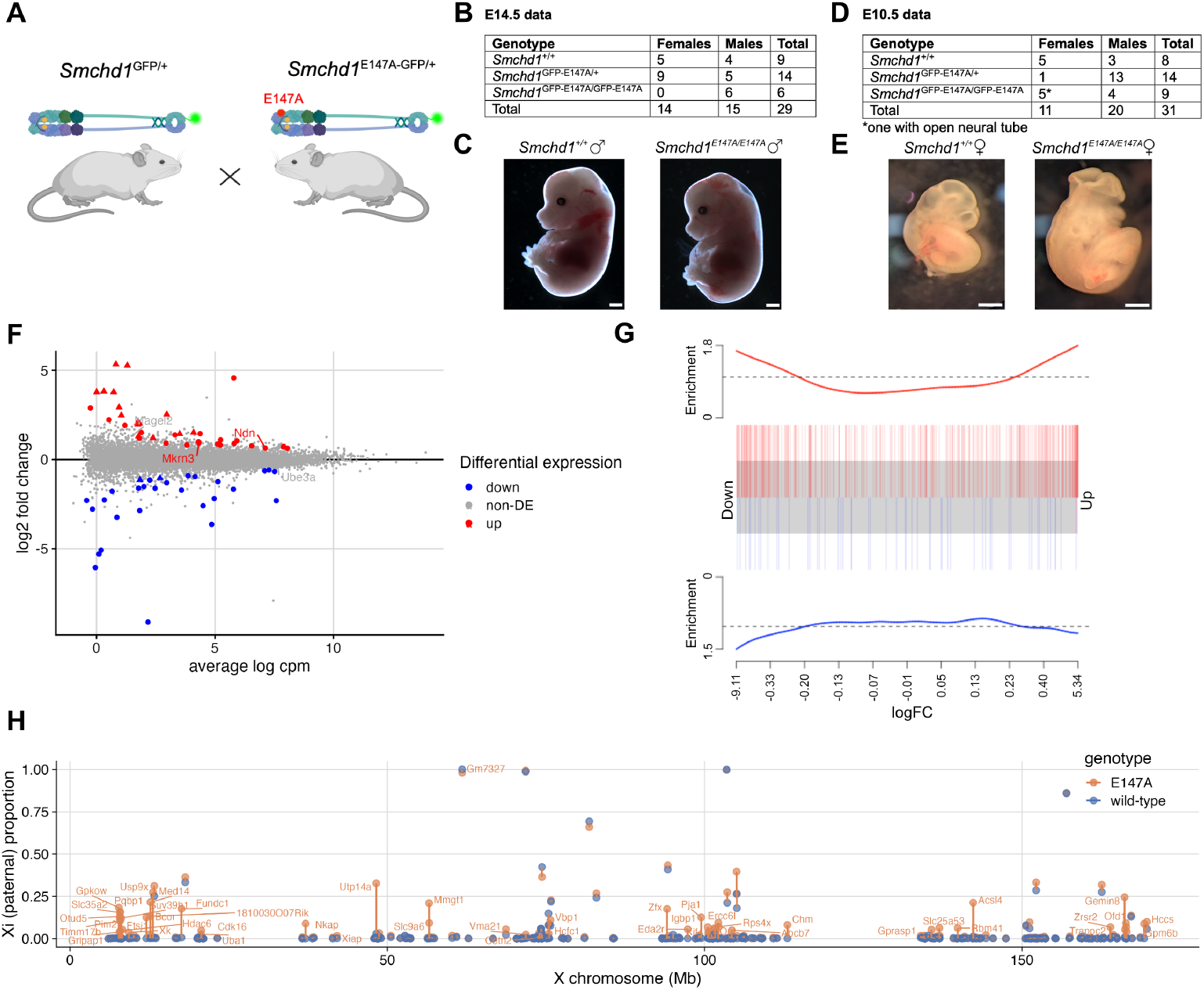
The E147A allele induces a *Smchd1*-null effect in a knockin mouse model. **(A)** A graphical representation of the *Smchd1*^GFP-E147A/+^ heterozygous intercrosses, showing **(B)** the resulting genotypes at E14.5 and **(D)** at E10.5 of development alongside **(C,E)** sample images of the corresponding embryos, where scale bars are 1 mm. **(F-H)** RNA-sequencing data from *Smchd1*^GFP-E147A/+^ and wild-type NSCs. **(F)** MA plot of average expression vs log2 fold change between the mutant and wild-type samples. Genes of interest are highlighted, where significantly differentially expressed genes (FDR<0.05) are red (upregulated) or blue (downregulated). Clustered protocadherins that are differentially expressed are marked with triangles. **(G)** Barcode plot comparing gene expression changes in *Smchd1*^GFP-E147A/+^ versus *Smchd1-*null data using Camera gene set test, p=0.0148. **(H)** Map of the X chromosome, showing the proportion of Xi-derived gene expression for each gene along the chromosome. The *Smchd1*^GFP-E147A/+^ data is shown in orange, and the wild-type data shown in blue. Differentially expressed genes (whose parental ratios differ between wild-type and mutant) are labelled.

We first examined the effect of the E147A mutation and the associated disrupted ATPase activity on embryonic viability. We performed *Smchd1*^GFP-E147A/+^ heterozygous intercrosses and dissected embryos at E10.5 and E14.5. We found that female *Smchd1*^GFP-E147A/GFP-E147A^ embryos die prior to mouse embryonic day 14.5 (E14.5) of development, whereas male *Smchd1*^GFP-E147A/GFP-E147A^ embryos survive and appear normal at this stage (χ^2^ test p<0.05 for F:M ratio of homozygotes, **Fig. 1B,C**). At E10.5, we identified live male and female *Smchd1*^GFP-E147A/GFP-E147A^ embryos, representing approximately 25% of all embryos, as expected based on Mendelian proportions (**Fig. 1D,E**). One of the female embryos had an open neural tube, which we previously observed in *Smchd1*-null embryos (37). Mid-gestation female-specific lethality and male survival are consistent with the time of lethality for female and male *Smchd1*-null embryos on a C57BL/6J background (38,39). These data suggest that ATPase activity is required for normal SMCHD1 function *in vivo*.

To test the effect of SMCHD1 ATPase disruption on gene expression, we created neural stem cells (NSCs) from *Smchd1*^GFP-E147A/+^ and wild-type E14.5 embryos and performed RNA sequencing (n=3 each for male heterozygous mutant and wild-type, n=2 for female heterozygous mutant and wild-type). We examined heterozygous NSC samples as homozygous females were non-viable prior to E14.5, and we have previously extensively characterised gene expression changes and SMCHD1 chromatin occupancy in NSCs. Differential gene expression analyses using the male and female NSCs combined revealed 67 significantly differentially expressed genes: 38 upregulated and 29 downregulated (FDR<0.05, **Fig. 1F, Supplementary Table 2**). The upregulated genes include 15 clustered protocadherins from the α and β clusters, and two *Snrpn* cluster imprinted genes (*Mkrn3* and *Ndn*) that are known SMCHD1 targets also upregulated in *Smchd1* null NSCs (11,16). Two protocadherin γ genes were among the downregulated genes, again as they are in *Smchd1* null NSCs (**Fig. 1F**, (11,16)). There was a significant overlap in observed differential expression between *Smchd1*^GFP-E147A/+^ and wild-type, and *Smchd1* null to wild-type NSCs (**Fig. 1G**, p=0.0148, gene set test), suggesting that disruption of even one allele of *Smchd1* via the E147A mutation is sufficient to cause a deficit in SMCHD1 silencing function.

The NSCs were produced to allow allele-specific analyses of X chromosome inactivation, as the paternal X chromosome is derived from the Castaneus strain, while the maternal X chromosome carries an *Xist*^deltaA^ allele on a 129-strain X chromosome (15). Given that Xist is required for silencing of the X chromosome *in cis*, the 129-derived X chromosome is always the active X chromosome and the Castaneus-derived X chromosome is the inactive X in this scenario. Of the 376 genes that had expressed SNPs with sufficient coverage, 50 showed a significantly altered proportion of paternal to maternal allele expression in the *Smchd1*^GFP-E147A/+^females compared with controls (FDR<0.05), 47 of which displayed increased expression from the (paternal) inactive X chromosome, indicative of compromised X inactivation (**Supplementary Table 3**). These genes are spread along the X chromosome (**Fig. 1H, Fig. S1**).

Taken together, the observed embryonic lethality of homozygous E147A mutant *Smchd1* and the gene expression alterations on autosomes and from the inactive X chromosome even in the heterozygote mutants, are consistent with SMCHD1’s ATPase activity being required for function in gene regulation *in vivo*.

### SMCHD1’s BAH and IGL-1 domains form a DNA-interaction site

FSHD-associated variants outside the core ATPase domain are known to compromise SMCHD1 ATPase activity (5,8,40). Hence, we sought to dissect the contribution of the two adjacent domains to SMCHD1 function using recombinant proteins. Based on their predicted structural homology with other functional domains, we termed these the Bromo-adjacent homology (BAH), as named previously (6), and immunoglobulin-like 1 (IGL-1) domains, comprising residues 571-688 and 688-819, respectively (**Fig. 2A**). Expression trials of the IGL-1 domain alone were unsuccessful, indicating poor protein stability. Following successful expression and purification of the SMCHD1 BAH domain alone (residues 571-688) and the BAH together with IGL-1 (residues 571-819) (**Fig. 2B,C**), we used a fluorescence polarization-based assay to examine whether these interact with nucleic acids as BAH domains are commonly found in chromatin-associated proteins (41,42). Indeed, we observed that BAH+IGL-1 was able to interact with both single- and double-stranded DNA (ssDNA and dsDNA), whereas the BAH alone or the extended SMCHD1 ATPase domain (residues 25-702), which incorporates the BAH domain, showed negligible binding, suggesting that DNA interaction requires the IGL-1 domain (**Fig. 2D,E**). Using varying lengths of either ssDNA or dsDNA probes, we estimated an affinity of ∼2 µM for ssDNA (**Fig. 2F**).

**Figure 2.**
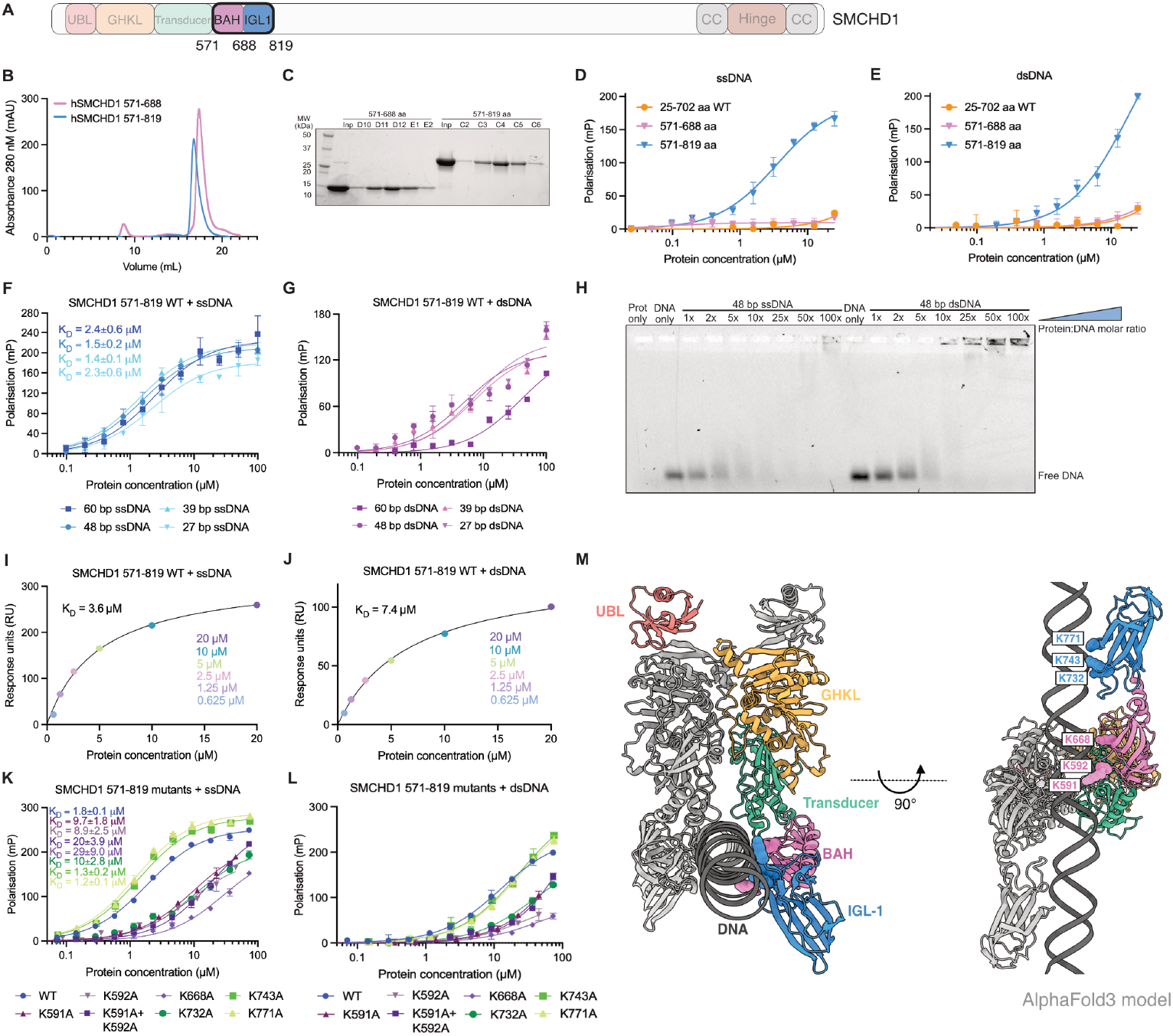
SMCHD1’s BAH and IGL-1 domains form a DNA-interaction site. (**A**) A schematic depicting the SMCHD1 domain architecture, highlighting the BAH and IGL-1 domains. (**B**) A size exclusion chromatography (SEC) profile of the BAH and BAH+IGL-1 domains, showing (**C**) a reducing SDS-PAGE gel of the eluting fractions. (**D-G**) Fluorescence polarization-based DNA-binding assays examining the binding of (**D-E**) various SMCHD1 constructs to a 6’FAM 48 base-pair (bp) (**D**) ssDNA or (**E**) dsDNA, and of the (**F-G**) SMCHD1 571-819 (BAH-IGL-1) domain binding to various lengths of 6’FAM-labelled (**F**) ssDNA or (**G**) dsDNA probes. Measurements were performed in duplicates and the graphs shown are representative of n=3 experiments for (**F)** and n=2 for (**G).** (**H**) EMSA examining the binding of wild-type 571-819 residue SMCHD1 to either 48 bp ssDNA or dsDNA with increasing protein: DNA molar ratios. (**I-J**) SPR analyses at steady state of SMCHD1 571-819 (BAH+IGL-1) with (**I**) 60 bp ssDNA or (**J**) 60 bp dsDNA, at varying protein concentrations. (**K-L**) Fluorescence polarization-based DNA-binding assays examining the affinity of various BAH-IGL-1 point mutants for 48 bp (**K**) ssDNA or (**L**) dsDNA probes. Measurements were performed in duplicates and the graphs shown are representative of n=2 experiments. (**M**) An AlphaFold3 model of the 25-819 amino acid SMCHD1 dimer in complex with double-stranded DNA with residues subjected to alanine mutagenesis highlighted.

Polarization values for dsDNA probes were consistently lower than observed for ssDNA and were not able to generate accurate K_D_ values (**Fig. 2G**), therefore we performed both electromobility shift assays (EMSA) and surface plasmon resonance (SPR) experiments as orthogonal methods to examine DNA binding. Both experiments suggested that SMCHD1’s BAH-IGL-1 domains exhibit an affinity in the low micromolar range for both ssDNA and dsDNA (**Fig. 2H-J**), highlighting a technical issue in the fluorescence polarization assay that specifically arises with the use of dsDNA. Using SPR in a kinetic configuration, we observed fast association and dissociation rates that indicate the interaction between SMCHD1’s BAH-IGL-1 and DNA occurs on a short timescale (**Fig. S1**). We used steady-state analysis to calculate affinity values, and obtained K_D_ values of 3.6 µM for ssDNA and 7.4 µM for dsDNA (**Fig. 2I,J**). Taken together, these results suggest that SMCHD1’s BAH and IGL-1 domains form a DNA-interaction site of a low micromolar affinity, within the same range previously observed for the SMCHD1 hinge domain, the first identified DNA-binding site in SMCHD1 (10,11).

To map the DNA-interaction site within the SMCHD1 BAH-IGL-1 module, we performed alanine mutagenesis of lysine residues that were predicted by an AlphaFold3 model (generated in complex with DNA) to be located within close proximity to the DNA probe. We targeted residues K591, K592 and K668 in the BAH domain and residues K732, K743 and K771 within the IGL-1 domain, and performed DNA-binding assays on the resulting SMCHD1 BAH-IGL-1 recombinant proteins to assess their affinity for nucleic acids, in comparison to the wild-type counterpart (**Fig. 2K,L**). All but two point mutations resulted in reduced affinity for DNA, with the most striking effect observed for the K668A mutant for which we obtained a K_D_ value of∼29 µM, compared to ∼1.8 µM obtained for wild-type SMCHD1 in the same assay (*p* <0.0001) (**Fig. 2K,L**). An additive effect was observed for the double mutant K591A+K592A, with an estimated K_D_ value of ∼20 µM, compared to K591A and K592A individually which displayed K_D_ values of ∼9.7 µM and ∼8.9 µM, respectively (*p* <0.0001). K743A and K771A variants surprisingly showed a slightly enhanced affinity for DNA, with K_D_ values of 1.3 µM and 1.2 µM (versus K_D_ ∼1.8 µM for WT SMCHD1, *p* <0.0001), suggesting these two lysine residues that are located within the IGL-1 domain may not have a direct role in nucleic acid interaction and their substitution may instead allow an easier entry for DNA.

### Cryo-EM of the wild-type SMCHD1 dimer (residues 25-819) reveals low resolution structures of the BAH and IGL-1 domains

SMCHD1’s BAH and IGL-1 domains are small proteins with molecular masses of ∼14 kDa each, which is below the molecular weight threshold that can be structurally interrogated with single-particle cryo-EM. To obtain a suitable protein sample, we instead produced recombinant SMCHD1 that encompasses the ATPase module as well as the adjacent BAH and IGL-1 domains (residues 25-819) (**Fig. 3A**). In the presence of the non-hydrolysable ATP (AMP-PNP), SMCHD1 ATPase-BAH-IGL-1 forms a stable dimeric complex with an approximate molecular weight of ∼180 kDa. After separating the monomeric and dimeric populations using size exclusion chromatography (SEC), fractions containing the dimeric protein complex were isolated and vitrified for cryo-EM analysis (**Fig. S2**). Three-dimensional reconstruction to an overall 3.1 Å resolution revealed the structure of the wild-type human SMCHD1 ATPase. While we were able to visualize the overall arrangement of the BAH and IGL-1 domains in the electron density map, the BAH domain was captured at an average resolution of ∼5 Å and the IGL-1 to approximately ∼7 Å which does not allow us to make detailed interpretations of these regions (**Fig. 3B**). We used an AlphaFold2 model of SMCHD1 ATPase-BAH-IGL-1 to refine against the electron density map, and obtained a structure that is very similar to the AlphaFold model itself (**Fig. 3C**).

**Figure 3.**
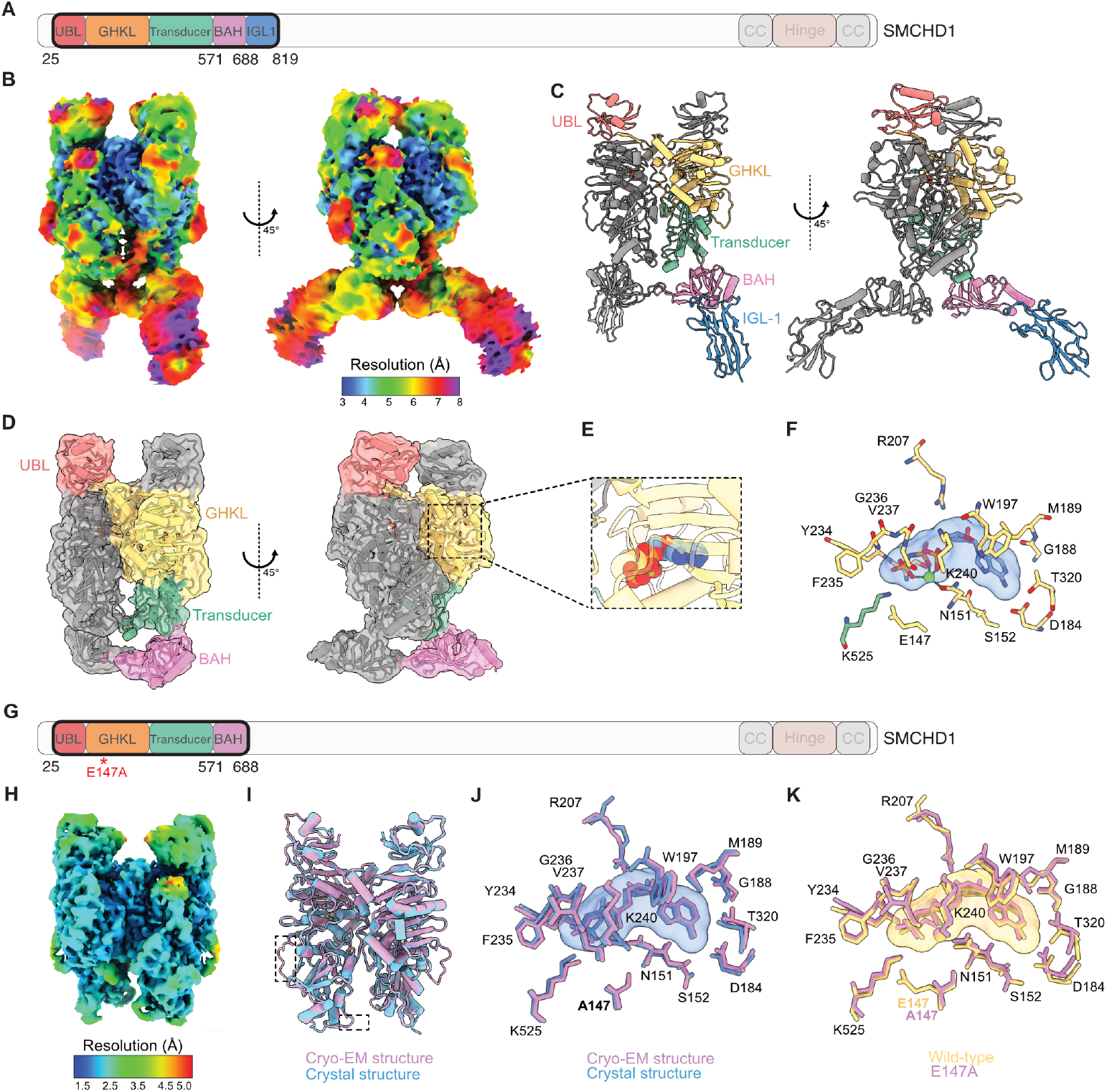
Cryo-EM of the wild-type SMCHD1 dimer (residues 25-819) reveals low resolution structures of the BAH and IGL-1 domains. **(A)** Schematic depicting the SMCHD1 domain architecture, highlighting the ATPase-BAH-IGL1 (residues 25-819) SMCHD1 construct. **(B)** A local resolution map of the cryo-EM density obtained, ranging between 3-8 Å (PDB:23DV, EMD-68889). **(C)** Modelled structure of the cryo-EM density map, obtained using an AlphaFold 3 model of the ATPase-BAH-IGL1 (residues 25-819) SMCHD1 protein. **(D)** The structure from **(B)** viewed with a high contour level, encompassing the ATPase-BAH (residues 25-688) of SMCHD1. **(E-F)** AMP-PNP-Mg^2+^ binding to the active site, highlighting **(F)** residues within 3 Å of the bound ligand (blue surface), with dashed lines representing coordination bonds with the Mg^2+^. **(G)** Schematic highlighting the domain architecture of SMCHD1, emphasizing the ATPase-BAH (residues 25-688) E147A SMCHD1 construct. **(H)** A local resolution map of the cryo-EM density obtained for the ATPase-BAH (residues 25-688) E147A SMCHD1 protein in complex with AMP-PNP/Mg^2+^, where density can only be observed for residues 25-580 (PDB:23DN, EMD-68884). **(I)** A final 3D structure of the 25-580 residue E147A SMCHD1 cryo-EM model (pink) superimposed with the previously solved crystal structure of 25-580 residue E147A SMCHD1, PDB:6MW7 (blue), highlighting two loop regions that are observed in our model (black dashed lines). **(J)** Superimposed active site residues from our E147A cryo-EM model (pink) and PDB:6MW7 (blue), and of **(K)** our E147A (pink) with our wild-type (yellow) cryo-EM model.

Similar to the previously published crystal structure of the E147A variant (8), each monomer within the SMCHD1 dimer adopts a head-to-head arrangement and comprises an N-terminal ubiquitin-like domain (UBL), a GHKL-type catalytic ATPase, a transducer domain, and a BAH domain, followed by an IGL-1 domain that is not resolved in the high contour model (**Fig. 3D**). The ATPase domain consists of a conserved Bergerat fold which is characteristic of GHKL family members (43,44). Within this domain, we can visualize the active site of wild-type SMCHD1 bound to AMP-PNP and Mg^2+^, corresponding to the nucleotide-binding site observed in the E147A variant (8) (**Fig. 3E,F**). The conserved residue E147 functions as the catalytic base, activating a water molecule for ATP hydrolysis, while N151 coordinates the Mg^2+^ ion required for catalysis (**Fig. 3F**). Notably, K525, located within the transducer domain, extends into the active site to form a hydrogen bond with the phosphate group of AMP-PNP. This residue has previously been described as a ‘switch’ lysine in GHKL ATPases, where it is proposed to sense nucleotide binding and regulate conformational transitions between monomeric and dimeric states (8,45).

We furthermore utilized cryo-EM to obtain a structure of the E147A point mutant in an ATPase-BAH (residues 25-688) context (**Fig. 3G**), which was solved at an overall 2.1 Å **(Fig. 3H**). However, the observed density for the BAH domains (residues 571-688) was still very poor, suggesting a high degree of flexibility (**Fig. S3**). Upon superimposition of this model with the 25-580 E147A SMCHD1 structure previously reported using X-ray crystallography (8), we can observe a highly similar arrangement, with just two additional loop regions present in our cryo-EM structure which are localized just downstream of the GHKL, to the transducer domain. One of these comprises of residues 256-373, and the other of residues 493-502 (**Fig. 3I**). Superimposition of active site residues likewise revealed a conserved structural arrangement between the E147A cryo-EM versus crystallography structures (**Fig. 3J**), whereas the only difference between our wild-type and E147A cryo-EM models was the mutation itself (**Fig. 3K**). These findings highlight that the E147A substitution does not grossly perturb ATPase architecture, and that the observed functional changes of this variant arise from impaired ATP turnover rather than structural destabilisation.

### The ATPase activity of full-length SMCHD1 is stimulated by DNA

To ask about the role of the DNA binding related to ATPase activity, we produced a near full-length recombinant mouse SMCHD1 protein sample comprising residues 25-1959. This construct excludes the C-terminal nuclear localization signal (NLS)(46) to prevent the protein from being imported into the nucleus of insect cells, our mammalian protein expression system (**Fig. 4A**). This ensures the almost-full-length recombinant SMCHD1 protein does not interact and co-purify with cellular DNA, which was an issue we had previously encountered when expressing the full-length mouse SMCHD1 (residues 1-2007). Both SEC (**Fig. 4B**) and mass photometry (**Fig. 4D**) analyses indicated a dimeric species, with an expected molecular weight of ∼430 kDa. Using the same fluorescence polarization-based DNA-binding assay, we obtained a K_D_ value of ∼208 nM for ssDNA binding (**Fig. 4E**), however the polarization signal for dsDNA (not shown) was very low which made it difficult to interpret. Therefore, we once again performed EMSA as an orthogonal method to better examine SMCHD1’s affinity for dsDNA, which indicated a similar affinity for both ssDNA and dsDNA (**Fig. 4F**).

**Figure 4.**
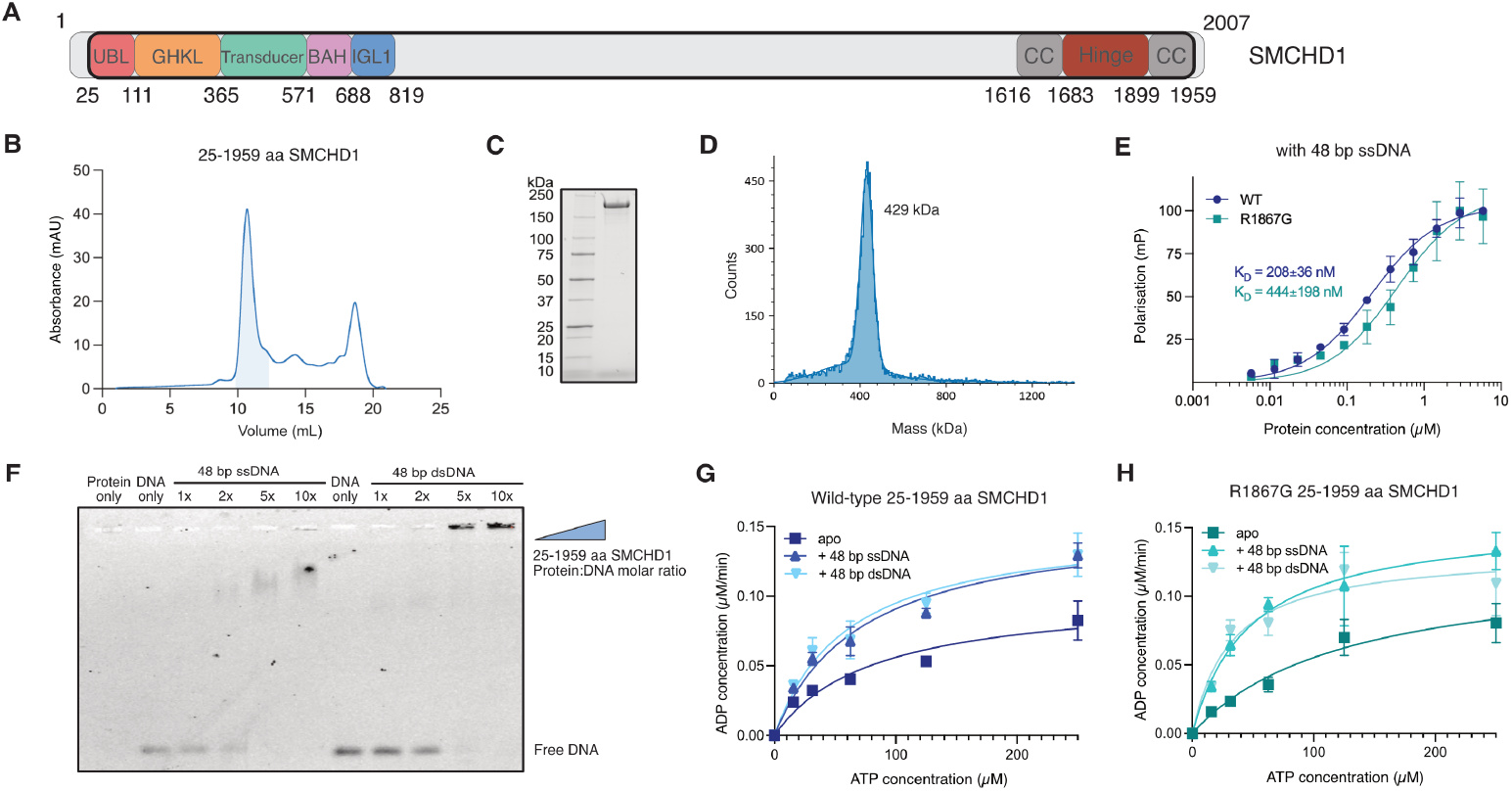
The ATPase activity of full-length SMCHD1 is stimulated by DNA. **(A)** A depiction of the gene architecture of SMCHD1, highlight the region of interest (residues 25-1959). **(B)** Size-exclusion chromatography (SEC) of the 25-1959 residue SMCHD1 protein, showing **(C)** a reducing SDS-PAGE gel and **(D)** mass photometry of the purified sample. **(E)** Fluorescence polarization-based assay examining the binding of wild-type and R1867G SMCHD1 protein to a 48-bp ssDNA probe (6’Fam labelled), where measurements were performed in duplicates and are representative of n=3 experiments. **(F)** Electromobility shift assay (EMSA) of the 25-1959 residue wild-type SMCHD1 with either 48-bp ssDNA or dsDNA, with the indicated molar ratios. **(G-H)** Fluorescence polarization ATPase assays examining the catalytic activity of **(G)** wild-type and **(H)** R1867G 25-1959 residue SMCHD1 in the presence and absence of 0.5 µM ssDNA or dsDNA. Measurements were performed in triplicates and the graphs are representative of n=2 experiments.

We simultaneously examined SMCHD1 point mutant R1867G, an FSHD-associated variant (R1866G in human SMCHD1) that displays a reduced affinity for DNA *in vitro* (9-11), in the context of the 25-1959 residue protein. Upon investigating its DNA-binding affinity using fluorescence polarization, we observed a K^D^ value of ∼444 nM for R1867G, compared to ∼208 nM obtained for WT SMCHD1 (*p* = 0.0005) (**Fig. 4E**). In our previous study, we investigated the R1867G variant in the context of the hinge domain only and we observed a ∼4-fold difference in DNA-binding affinity (K^D^ of 6.2 µM for R1867G vs 1.5 µM for WT) (10). This may be attributed to the presence of more nucleic acid-interacting residues in the full-length SMCHD1 protein compared to the hinge domain only, which can better compensate for the R1867G-induced decreased affinity for DNA.

We subsequently interrogated the *in vitro* ATPase function of these two protein samples, either in the presence or absence of ssDNA or dsDNA. Interestingly, we observed a ∼2-fold stimulation of ATP turnover in the presence of either ssDNA or dsDNA for wild-type SMCHD1, as well as for the R1867G point variant (**Fig. 4G-H**). DNA-stimulation of ATPase activity has not been previously reported for SMCHD1, and hence raises further questions about the underlying mechanistic details. These results also highlight that, despite the 2-fold decrease in DNA-binding affinity, the catalytic activity of the R1867G variant can be stimulated by DNA to the same degree as wild-type SMCHD1, further suggesting that the nucleic acid-induced stimulation of ATPase activity may be due to DNA binding to other sites of the SMCHD1 protein, such as the BAH-IGL1 domain.

### DNA-binding at SMCHD1’s BAH-IGL1 domains leads to ATPase stimulation

To determine if binding of DNA at the BAH-IGL1 region results in the observed stimulation of SMCHD1’s ATPase, we examined the 25-819 residue SMCHD1 construct that contains both the ATPase and the downstream BAH-IGL1 (**Fig. 5A**). First, we assessed its affinity for ssDNA *in vitro*, and obtained a K_D_ value of 3.9 µM compared to 1.8 µM observed for the BAH-IGL1 construct (residues 571-819), which we attribute to possible steric hindrance in the extended protein sample (**Fig. 5B**). Upon investigating the protein’s catalytic rate in the presence and absence of DNA, we found no significant difference in the rate of ATP turnover, suggesting it may not be the interaction of DNA at the BAH-IGL1 that leads to DNA-driven stimulation in this context (**Fig. 5C**).

**Figure 5.**
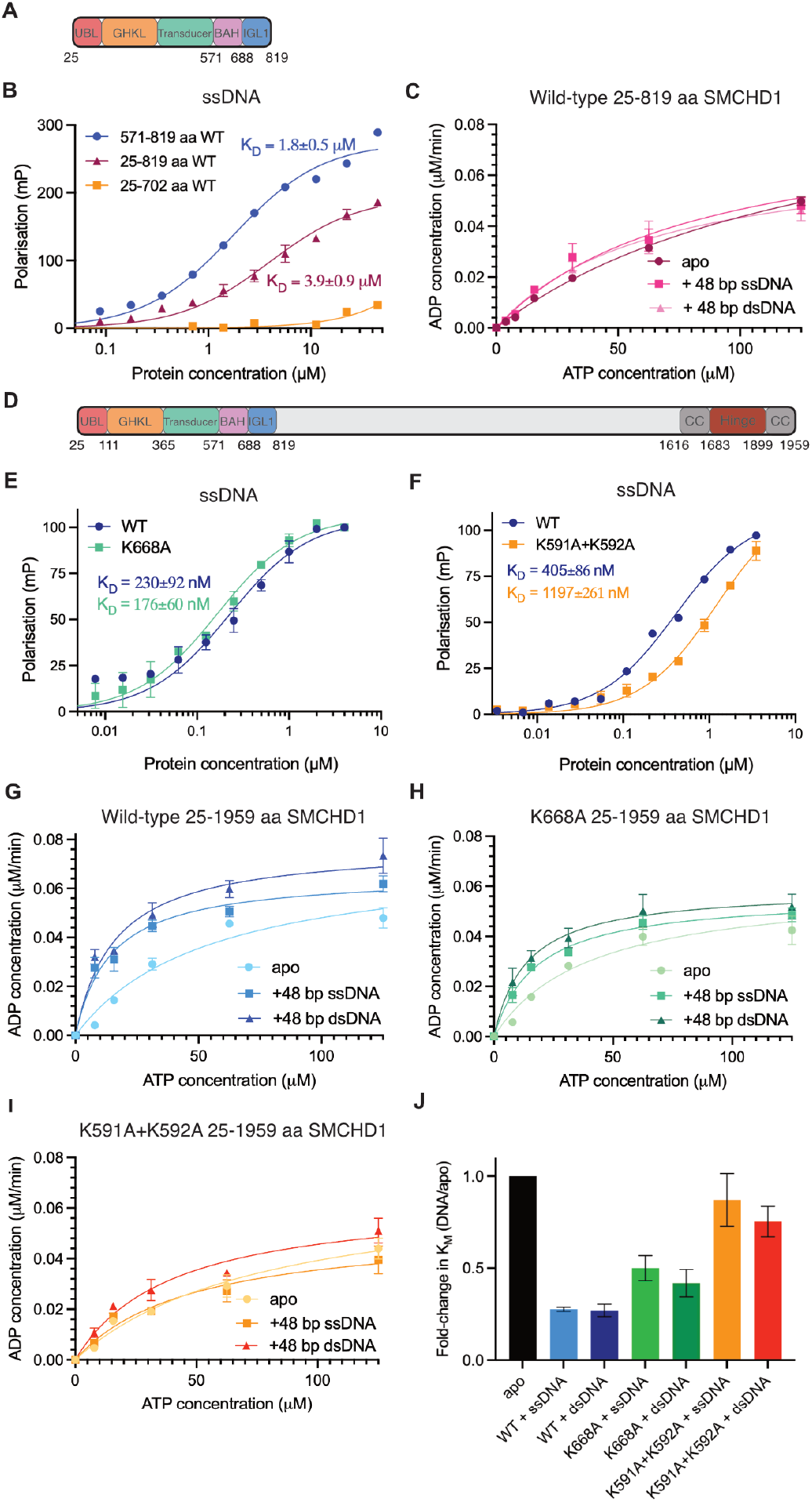
DNA-binding at SMCHD1’s BAH-IGL1 domains leads to ATPase stimulation. **(A)** A schematic depicting the gene architecture of the SMCHD1 ATPase-BAH-IGL1 construct (residues 25-819). **(B)** Fluorescence polarization-based DNA-binding assay using a 6’FAM 48 base-pair (bp) ssDNA with the depicted SMCHD1 constructs, where measurements were performed in duplicate and the graph is representative of n=3 independent experiments. **(C)** Fluorescence polarization ATPase assays examining the catalytic activity of wild-type 25-819 residue SMCHD1 in the presence and absence of 0.5 µM ssDNA or dsDNA. Measurements were performed in triplicate and the graphs are representative of n=2 experiments. **(D)** The gene architecture of the SMCHD1 construct encompassing residues 25-1959. **(E-F)** Fluorescence polarization-based DNA-binding assays using a 6-FAM 48 base-pair (bp) ssDNA with the depicted point variants of the 25-1959 residue SMCHD1 samples, where measurements were performed in duplicate and the graph is representative of n=3 independent experiments. **(G-I**) Fluorescence polarization ATPase assays examining the catalytic activity of **(G)** wild-type, **(H)** K668A and **(I)** K591A+K592A 25-1959 residue SMCHD1 in the presence and absence of 0.5 µM ssDNA or dsDNA. **(H)** A graphical representation of the fold-change in K_M_ between the wild-type and mutant protein samples, in the presence or absence of DNA. Measurements were performed in triplicate and the graphs are representative of n=3 experiments for (**H)** and n=2 for (**I)**.

We further addressed this question in the context of the full-length SMCHD1 protein, which is most representative of its native function (**Fig. 5D**). To do so, we introduced point mutations K591A+K592A or K668A in the 25-1959 residue SMCHD1 construct, for which we previously observed a ∼10-fold and ∼15-fold decrease in DNA-binding affinity, respectively, using the SMCHD1 BAH-IGL-1 construct (**Fig. 2K-L**). Surprisingly, the *in vitro* binding assay showed no significant difference in the DNA-binding affinity of point mutant K668A (*p* = 0.0751) (**Fig. 5E**), suggesting possible compensation by other DNA-binding residues within the full-length SMCHD1 protein. However, we did observe a ∼3-fold reduction in DNA-binding affinity for the K591A+K592A double mutant (*p* <0.0001) (**Fig. 5F**).

When we further analysed the catalytic activity of these two SMCHD1 variants in the absence and presence of DNA, we observed a decreased DNA-induced stimulation of ATPase function in both when compared to the wild-type counterpart (**Fig. 5G-I**). Upon comparing the ATP affinity (K_m_) values for the wild-type and mutant samples in the presence and absence of DNA, we found that binding of either ssDNA and dsDNA reduces the respective K_m_ values and therefore enhance SMCHD1’s affinity for ATP. In the wild-type protein, the reduction in K_m_ was ∼4-fold compared to only ∼2-fold for the K668A variant (**Fig. 5J**). This effect was even less pronounced for the K591A+K592A mutant, as we observed no reduction in K_m_ in the presence of ssDNA (*p* = 0.2592), and only a ∼1.3-fold reduction in the presence of dsDNA (*p* = 0.0356) (**Fig. 5J**). These results suggest that DNA-binding at residues K668, and in particular K591/K592, play a key role in the observed enhanced affinity for ATP, and therefore that nucleic acid interaction at the BAH-IGL1 domain leads to stimulated ATP turnover in SMCHD1.

### Disruptive mutations in the BAH-IGL1 domains lead to an altered nuclear localization pattern of SMCHD1 in cells

We next sought to investigate the effects of mutations K591A+K592A and K668A (BAH domain) compared to E147A (ATPase domain) and corresponding controls, in a cellular context. We used the CRISPR/Cas9 system to introduce these mutations into human embryonic kidney (HEK) 293T cells, from which we generated clonal cell lines and confirmed the requisite base changes required to generate the specific amino acid substitutions were present on the *SMCHD1* alleles using Illumina sequencing.

Using immunofluorescence microscopy, we observed differences between the nuclear localisation pattern of all three mutant SMCHD1 cell lines relative to their corresponding ‘no guide RNA’ controls. In particular, each mutation disrupted SMCHD1’s localisation to the Xi, which is marked by trimethylation of Lys27 on Histone 3 (H3K27me3), with SMCHD1 instead diffusely distributed throughout the nucleus (**Fig. 6A**). Quantification of mean SMCHD1 intensity revealed reductions in both nuclear regions (**Fig. 6B,D,F**) and Xi foci (**Fig. 6C,E,G**). Loss of SMCHD1 at the Xi also corresponds to an increased H3K27me3 signal, consistent with previous studies that have shown increased spreading of H3K27me3 domains to regions not normally decorated by this mark, in the absence of SMCHD1 (**Fig. B-G**) (16,47). Taken together, these results suggest that both impaired ATP hydrolysis and/or DNA-binding via the BAH-IGL1 domain disrupt SMCHD1’s ability to maintain stable interactions at native binding sites on chromatin.

**Figure 6.**
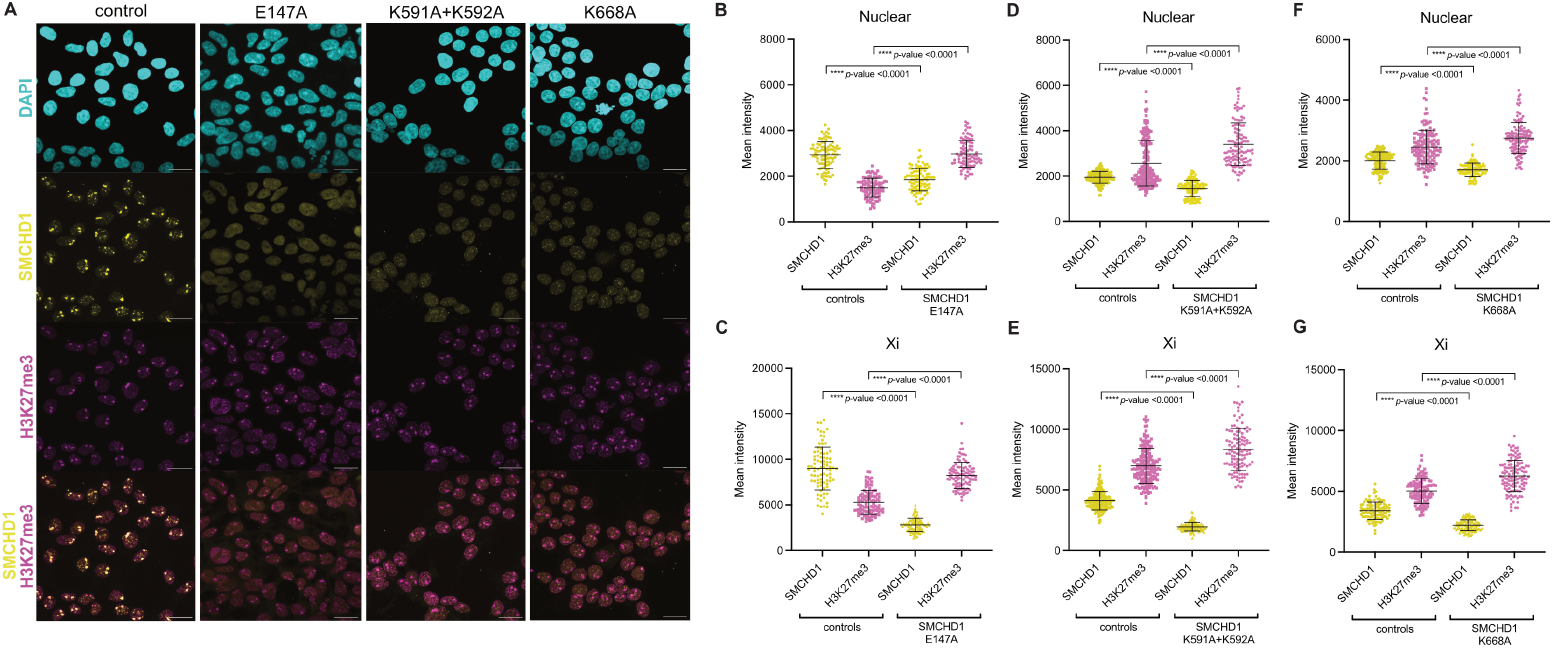
Disruptive mutations in the BAH-IGL1 domains lead to an altered nuclear localization pattern of SMCHD1 in cells. **(A)** Immunofluorescence microscopy of a control cell line (no guide RNA) that is representative of wild-type HEK293T cells, compared to CRISPR/Cas9-modified HEK293Ts containing point mutations: E147A, K591A+K592A and K668A. DAPI staining is shown in cyan, SMCHD1 in yellow and H3K27me3 in magenta, with merged images shown below as indicated. Maximum intensity projection images are shown as representative of n>100 cells per cell line. Scale bars, 20 µm. **(B-E)** Mean intensity values of SMCHD1 and H3K27me3 in **(B-C)** E147A, **(D-E)** K591A+K592A and **(F-G)** K668A HEK293Ts and their respective controls, for either the whole nucleus or just the Xi.

## DISCUSSION

In this study, we used genetic, biochemical and structural approaches to dissect how ATP hydrolysis and DNA binding contribute to SMCHD1 function. Using our mouse model, we found that the *Smchd1*^GFP-E147A^ allele resembles a *Smchd1*-null allele, where absence of SMCHD1 leads to female-specific embryonic lethality due to the failure of X-chromosome inactivation. These findings indicate that ATP hydrolysis is required for normal SMCHD1 function *in vivo*, likely for its stable association with chromatin at target loci. Consistent with these results, heterozygous FSHD-associated mutations in *SMCHD1* that result in a complete loss of catalytic function have been identified exclusively in male patients (**Supplementary Table 4**), providing human evidence for female-specific incompatibility arising from impaired X-chromosome inactivation even due to only one *SMCHD1* allele being affected.

We next identified the BAH and IGL-1 domains as a novel DNA-interaction site that is located immediately C-terminal to SMCHD1’s ATPase module. Alanine mutagenesis of conserved lysine residues revealed that DNA binding is distributed across both domains, with some residues contributing additively to the interaction. The presence of multiple DNA-binding sites within SMCHD1 suggests that chromatin engagement is likely multivalent, potentially enabling SMCHD1 to bridge or stabilise higher-order chromatin structures. Such interactions may be particularly important for SMCHD1’s role regulating chromatin architecture at large repressed domains, including the Xi and clustered gene loci such as the *Hox* clusters and the clustered protocadherin genes (11,16,47-49). Further supporting this idea, recent *in vitro* single-molecule studies suggested that SMCHD1 forms DNA clusters by bringing together different segments of DNA, leading to compaction (50).

Our cryo-EM analysis of the SMCHD1 ATPase-BAH-IGL1 dimer (residues 25-819) provides the first structural insight of the wild-type human ATPase in complex with AMP-PNP/Mg^2+^, while also revealing the overall arrangement of the downstream BAH and IGL-1 domains. Although these domains were resolved at lower resolution, their apparent flexibility relative to the ATPase core may be due to the absence of the C-terminal hinge domain and surrounding coiled-coils which represent a constitutive dimerization site of two SMCHD1 protomers and may strengthen the structural integrity of the overall dimer (2,10). Further work will focus on obtaining a cryo-EM structure of the aforementioned extended ATPase dimer in complex with DNA, which may rigidify the BAH and IGL-1 domains by providing a bridging site in the absence of the C-terminal hinge domain. While structural studies of the full-length SMCHD1 protein are also a key objective, our previous attempts and published negative staining EM data have indicated that the conformational heterogeneity of the rod-shape dimer presents a challenge for structural determination (6).

A key finding of this study is that DNA stimulates the ATPase activity of full-length SMCHD1. Specifically, we found that DNA binding at the BAH-IGL-1 site enhances SMCHD1’s affinity for ATP *in vitro*, suggesting that this interaction with DNA promotes a conformational change in the SMCHD1 protein that favours ATP-binding. This property has not been previously described for SMCHD1 and proposes a direct coupling between chromatin engagement and catalytic activity. While DNA was not present in our cryo-EM structure, we hypothesise that DNA-binding at the BAH-IGL-1 site likely brings the two SMCHD1 ATPases within the dimer in closer proximity, with one DNA strand bridging the two DNA-binding sites, as predicted by AlphaFold3 (**Fig. 2M**). Such a mechanism would augment the rate of ATP turnover by reducing the energetic requirement of cycling between the monomer/dimer conformations.

Interestingly, FSHD-associated *SMCHD1* missense variants within the BAH-IGL-1 region (W596G, V615D, P622L, V641L, P690S, L748P and Y774C) (2) do not map to residues that are predicted to interact with DNA. In addition, only one reported variant within the SMCHD1 hinge domain, R1867G, showed reduced affinity for DNA *in vitro*, although did not markedly reduce DNA affinity compared to wild-type SMCHD1 in the context of full-length protein in the current study. This may suggest that, in FSHD patients that express *SMCHD1*, the protein’s ability to interact with nucleic acids is largely preserved, highlighting the potential for exploiting the observed DNA-induced stimulation of SMCHD1’s ATPase activity as a therapeutic strategy for FSHD.

Our cellular findings are consistent with recent single-molecule data demonstrating that the SMCHD1 E147A variant, which impairs ATP hydrolysis, alters chromatin-binding specificity in cells (12). Although this variant exhibits a substantial chromatin-bound population within the nucleus relative to wild-type cells, it fails to show enrichment at the inactive X chromosome, indicating that ATP hydrolysis is not required for chromatin association but for binding specificity at target loci. In agreement with this model, we find that mutations disrupting either ATP hydrolysis (E147A) or DNA binding via the BAH–IGL-1 module lead to loss of stable SMCHD1 localisation at the inactive X chromosome and instead display a diffuse nuclear distribution. These defects are accompanied by increasing spreading of H3K27me3 marks (16,47), consistent with compromised silencing at the Xi. Together, these observations support a model in which SMCHD1 requires both ATP-driven conformational cycling and bidentate DNA binding to achieve stable chromatin residency *in vivo*.

Based on our findings, we propose that SMCHD1 binds chromatin through at least two DNA-interaction sites—the C-terminal hinge domain and the BAH-IGL-1 module adjacent to the N-terminal ATPase. DNA engagement at the BAH-IGL-1 stimulates ATP hydrolysis by likely bringing SMCHD1’s ATPase modules in closer proximity, which upon ATP-binding leads to homodimerization. By additionally providing a second DNA-binding site, this allows SMCHD1 to bring together different chromatin segments that can alter chromatin architecture (16,48,51) and may ultimately lead to compaction (50). Precisely how SMCHD1 localizes to target loci within the genome to prompt specific gene expression changes remains an open question, although it occurs downstream of H2AK119ub at least for the inactive X chromosome (13,51). We propose that disruption of ATP hydrolysis, such as by the introduction of the E147A mutation, prevents conformational cycling between the monomeric and dimeric states of the ATPase modules (8) and uncouples chromatin binding from stable residency, leading to impaired X-chromosome inactivation and transcriptional de-repression. Such a mechanism would align with SMCHD1’s proposed role as a chromatin architectural protein rather than a classical motor enzyme (50), where its inherently slow GHKL-type ATPase activity (5-7) likely only generates sufficient energy for local conformational changes.

Future studies will be required to determine how these mechanisms operate in a cellular context, and whether other factors such as the proposed interacting protein LRIF1 (12) are also able to modulate SMCHD1’s ATPase activity *in vivo*. Nevertheless, our work establishes ATP hydrolysis and bidentate DNA binding as critical features of SMCHD1 function and provides a framework for understanding how SMCHD1 mutations disrupt gene regulation in development and disease.

## DATA AVAILABILITY

CryoEM protein structures were deposited in the PDB/EMDB: 25-688 residue E147A SMCHD1, PDB: 23DN, EMD-68884; 25-819 residue wild-type SMCHD1, PDB:23DV, EMD-68889. The RNA sequencing data have been uploaded to the Gene Expression Omnibus, and are available under accession GSE316900.

## SUPPLEMENTARY DATA

Supplementary Data are available at *NAR* Online.

## ACKNOWLEDGEMENTS

We acknowledge the WEHI Protein Production Facility (PPF) for the production of various SMCHD1 protein samples used in this study. We thank the Melbourne Advanced Genome Editing Centre (MAGEC) Laboratory for their assistance with the generation of the *Smchd1*^GFP-E147A^ mouse. We thank the electron microscopy facility and their staff at the Ian Holmes Imaging Centre (IHIC), located at the Bio21 Molecular Science and Technology Institute.

## AUTHOR CONTRIBTUTIONS

A.D.G conceptualised the experiments, analysed data, acquired funding, performed investigations, validated and visualised data and wrote the draft manuscript. R.W.B. analysed data, acquired funding, performed investigations and validated data. A.L. performed investigations and wrote the draft manuscript. T.C. performed investigations and visualised data. M.I., K.B., I.W. and S.E. performed investigations. H.K.V. conceptualised the experiments, acquired funding and supervised the work. Q.A.G. analysed data, acquired funding, visualised data and wrote the draft manuscript. P.E.C. acquired funding and supervised the work. J.M.M. conceptualised the experiments, acquired funding, supervised the work and validated the data. M.E.B. conceptualised the experiments, acquired funding, supervised and validated the work, and drafted the manuscript. All authors edited the manuscript.

## FUNDING

This work was supported by an FSHD Society Fellowship [5059384059] and Jack Brockhoff Early Career Grant [to A.D.G], Al and Val Rosenstrauss Fellowship from the Rebecca L. Cooper Medical Research Foundation [F20221078 to H.K.V.], National Health and Medical Research Council of Australia (NHMRC) Investigator grants [2007996 to Q.A.G.; 2016894 to R.W.B; 2009062 to P.E.C.; 1172929 and 2034104 to J.M.M.; 1194345 and 2041117 to M.E.B.], with NHMRC IRIISS and Victorian State Government Operational Infrastructure Support.

## Notes

### Competing Interest Statement

The authors have declared no competing interest.

