## Supplementary Figures for "DNA binding by ATPase-adjacent domains stimulates SMCHD1 ATPase activity"

### Supplementary Figure 1

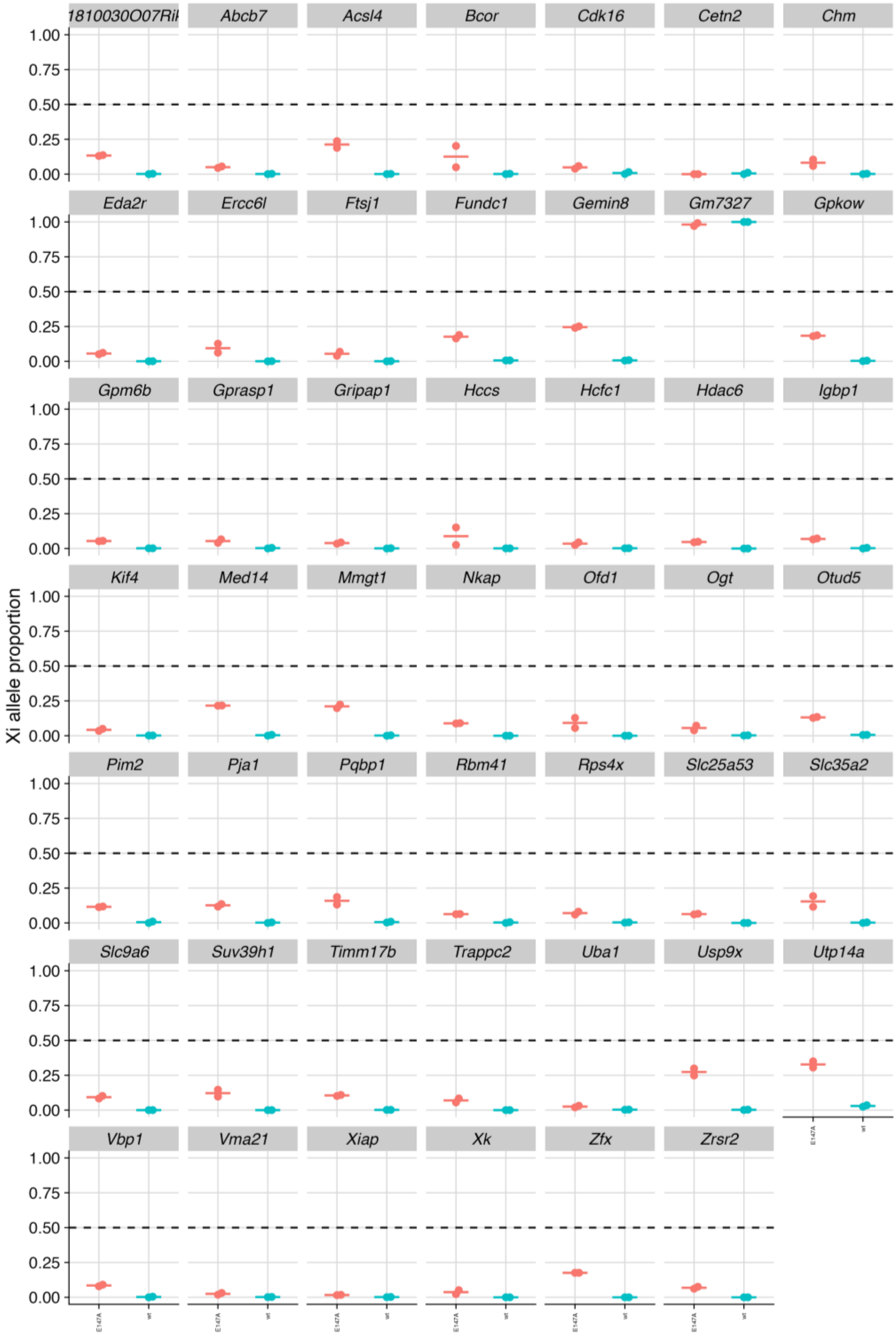


**Figure S1. Upregulation of the inactive X chromosome in *Smchd1*$^{E147A\text{-}GFP/+}$ NSCs**
Allele-specific RNAseq data shown for *Smchd1*$^{E147A\text{-}GFP/+}$ (red, n=2) and *Smchd1*$^{+/+}$ (teal, n=2) NSC samples. All genes shown are statistically significantly increased in the *Smchd1*$^{E147A\text{-}GFP/+}$ samples (FDR<0.05). Expression levels of the inactive X chromosome are shown as a proportion of the total expression for each gene.

### Supplementary Figure 2

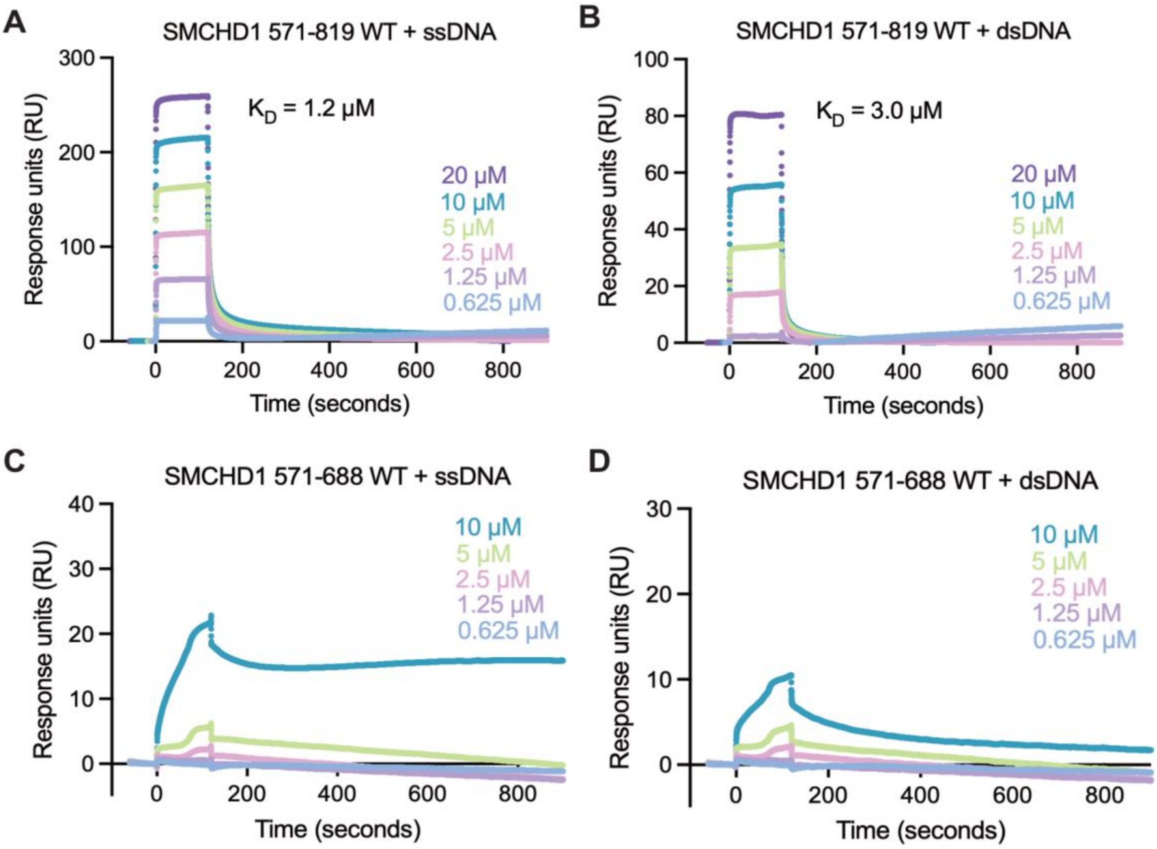


**Figure S2. Surface plasmon resonance (SPR) in a kinetic configuration.** SPR of wild-type SMCHD1 residues 571-819 (BAH+IGL1) with (A) 60 bp single-stranded DNA or (B) 60 bp double-stranded DNA, and of wild-type SMCHD1 residues 571-688 (BAH) with (C) 60 bp single-stranded DNA or (D) 60 bp double-stranded DNA. Varying protein concentrations used are depicted in different colours, as shown above.

### Supplementary Figure 3

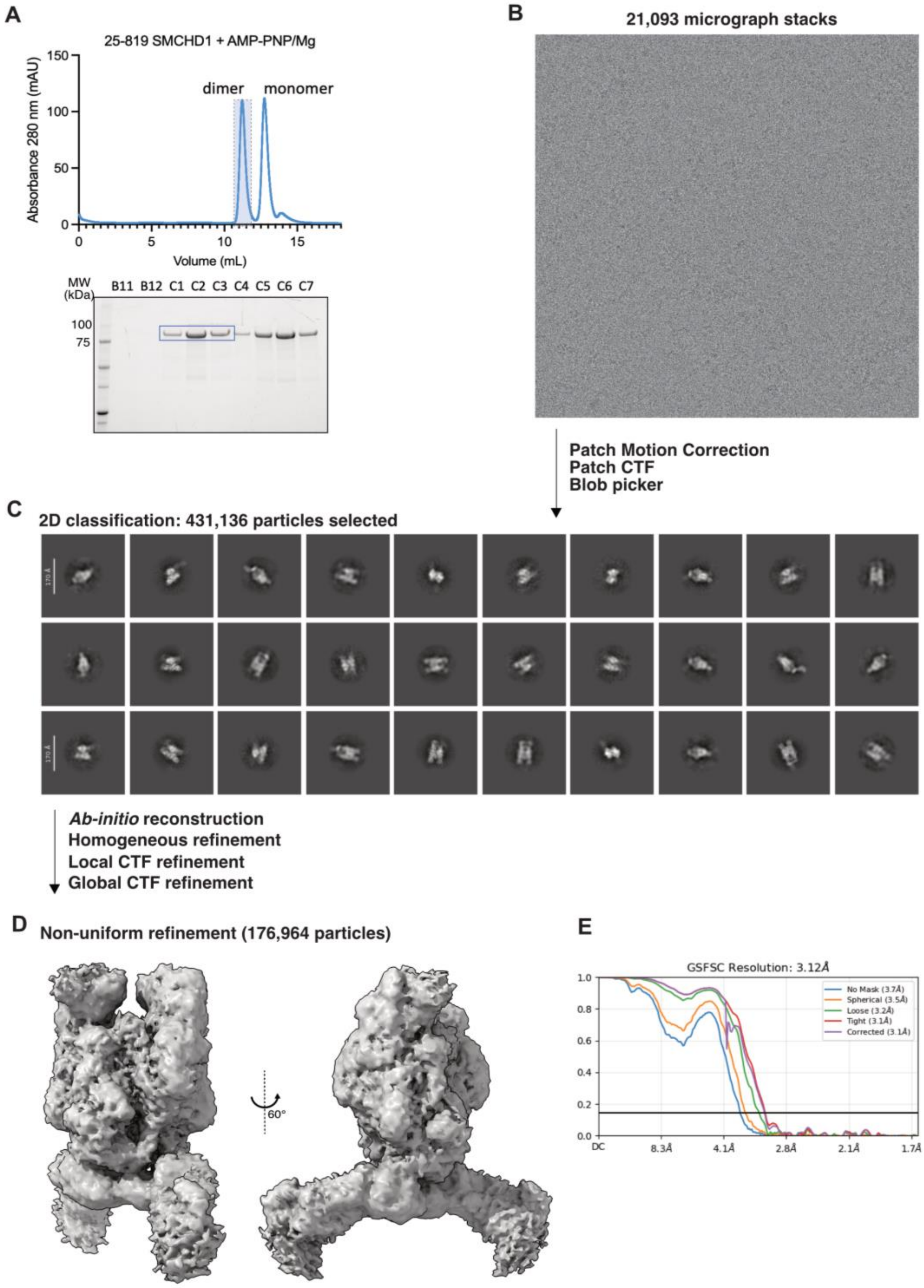


**Figure S3. Cryo-EM data acquisition and data processing. (A)** A size exclusion chromatography profile of the wild-type 25-819 residue SMCHD1 in the presence of non-hydrolysable AMP-PNP/$Mg^{2+}$, highlighting the dimeric peak used for this cryo-EM dataset. **(B)** We obtained 21,093 micrographs which were processed in CryoSPARC as briefly described above. **(C)** Particles were selected from 2D classification, followed by several refinement steps and a **(D)** final non-uniform refinement model obtained from 176,964 particles, with an **(E)** overall resolution of 3.12 Å.

### Supplementary Figure 4

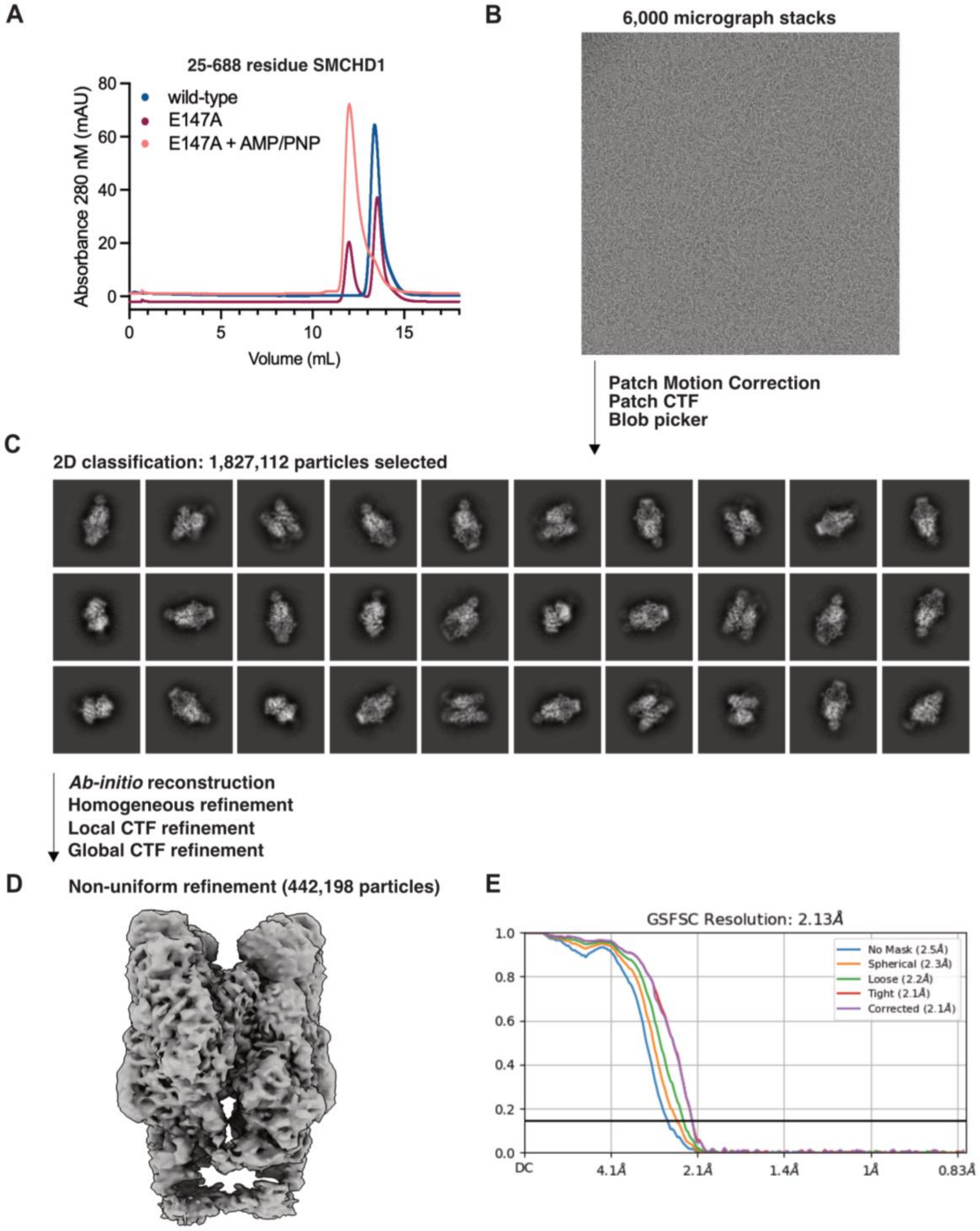


**Figure S4. Cryo-EM data acquisition and data processing. (A)** A size exclusion chromatography profile of the wild-type and E147A 25-688 residue SMCHD1, in the absence and presence of non-hydrolysable AMP-PNP/$Mg^{2+}$. For this cryo-EM dataset, we collected the 25-688 E147A + AMP-PNP/$Mg^{2+}$ dimer peak (orange line). **(B)** We obtained 6,000 micrographs which were processed in CryoSPARC as briefly described above. **(C)** Particles were selected from 2D classification, followed by several refinement steps and a **(D)** final non-uniform refinement model obtained from 442,198 particles, with an **(E)** overall resolution of 2.13 Å.
