## Supplementary Table 1 for "DNA binding by ATPase-adjacent domains stimulates SMCHD1 ATPase activity"

| Reagent | Purpose | Sequence |
|---|---|---|
| Smchd1 E147A gRNA 1 | Creation Smchd1 E147A mice | GATGGTTTGAATGACCCCAC |
| Smchd1 E147A gRNA 2 | Creation Smchd1 E147A mice | GGCGGGAAAGCAATCTGTTT |
| Smchd1 E147A oligo F | Amplify around Smchd1 E147A to genotype | ACGTGGTTGCTTTAAGGTGC |
| Smchd1 E147A oligo R | Amplify around Smchd1 E147A to genotype | GTAAAATATTCAAGTGCTTACAGTGC |
| Smchd1 F | Allelic discrimination genotyping assay | GGAGAGGTTAAGCACTGTTTTTCAA |
| Smchd1 R | Allelic discrimination genotyping assay | ACCATTATTTCGAGAAGTAGCTGACAA |
| Smchd1 E147A probe | Allelic discrimination genotyping assay | CTTGCAGAGTTAATCG |
| Smchd1 WT probe | Allelic discrimination genotyping assay | TTGCAGCCTTAATCG |
| SMCHD1 K591A F | Mutagenesis | GATCTTCCTTCTGCAAAGCAAGGTCCCTGG |
| SMCHD1 K591A R | Mutagenesis | CCAGGGACCTTGCTTTGCAGAAGGAAGATC |
| SMCHD1 K592A F | Mutagenesis | GATCTTCCTTCTAAAGCGCAAGGTCCCTGG |
| SMCHD1 K592A R | Mutagenesis | CCAGGGACCTTGCGCTTTAGAAGGAAGATC |
| SMCHD1 K591A+K592A F | Mutagenesis | CCTGATCTTCCTTCTGCAGCGCAAGGTCCCTGG |
| SMCHD1 K591A+K592A R | Mutagenesis | CCAGGGACCTTGCGCTGCAGAAGGAAGATCAGG |
| SMCHD1 K668A F | Mutagenesis | GTGCCAATTGCAGCGCTGGATAGGACAG |
| SMCHD1 K668A R | Mutagenesis | CTGTCCTATCCAGCGCTGCAATTGGCAC |
| SMCHD1 K732A F | Mutagenesis | GAAGCAATGCAAGCGCTTCCAGGAACAAG |
| SMCHD1 K732A R | Mutagenesis | CTTGTTCCTGGAAGCGCTTGCATTGCTTC |
| SMCHD1 K743A F | Mutagenesis | GGAGGGTCAAAGGCACTCCTGGTTGAG |
| SMCHD1 K743A R | Mutagenesis | CTCAACCAGGAGTGCCTTTGACCCTCC |
| SMCHD1 K771A F | Mutagenesis | CAACATGGAGGAGCATGGCCTTACTGG |
| SMCHD1 K771A R | Mutagenesis | CCAGTAAGGCCATGCTCCTCCATGTTG |
| Biotinylated 60 bp ssDNA/dsDNA | Surface plasmon resonance | 5`biotin-GGGTGAACCTGCAGGTGGGCAAAGATGTCCTAGCAAGGCACT GGTAGAATTCGGCAGCGT<br><br>ACGCTGCCGAATTCTACCAGTGCCTTGCTAGGACATCTTTGCCCACCT GCAGGTTCACCC |
| 6-Fam 5'labelled 60 | Fluorescence polarization | /56-FAM/ACGCTGCCGAATTCTACCAGTGCCTTGCTAGGACATCTTTGCCCACCT GCAGGTTCACCC<br><br>GGGTGAACCTGCAGGTGGGCAAAGATGTCCTAGCAAGGCACTGGTAGAATTC |

| | | |
|---|---|---|
| bp ssDNA/dsDNA | DNA-binding assay; EMSA | GGCAGCGT |
| 6-Fam 5'labelled 48 bp ssDNA/dsDNA | Fluorescence polarization DNA-binding assay; EMSA | /56-FAM/ACGCTGCCGAATTCTACCAGTGCCTTGCTAGGACATCTTTGCCCACCT<br><br>AGGTGGGCAAAGATGTCCTAGCAAGGCACTGGTAGAATTCGGCAGCGT |
| 6-Fam 5'labelled 39 bp ssDNA/dsDNA | Fluorescence polarization DNA-binding assay | /56-FAM/ACGCTGCCGAATTCTACCAGTGCCTTGCTAGGACATCTT<br><br>AAGATGTCCTAGCAAGGCACTGGTAGAATTCGGCAGCGT |
| 6-Fam 5'labelled 33 bp ssDNA/dsDNA | Fluorescence polarization DNA-binding assay | /56-FAM/ACGCTGCCGAATTCTACCAGTGCCTTGCTAGGA<br><br>TCCTAGCAAGGCACTGGTAGAATTCGGCAGCGT |
| 6-Fam 5'labelled 27 bp ssDNA/dsDNA | Fluorescence polarization DNA-binding assay | /56-FAM/ACGCT CCGAATTCTACCAGTGCCTTG<br><br>CAAGGCACTGGTAGAATTCGGCAGCGT |
| Human SMCHD1 E147A HDR donor oligo | CRISPR/cas9 mutagenesis of SMCHD1 in HEK293Ts | AGTTAAGCAATGTTTTCTTGATCTCTTGCAGCATTTGCTCTGGCCGCATTAATTGACAATTCATTGTCTG CTACTTCTCGTAACATT |
| Human SMCHD1 K591A+K592A crRNA | CRISPR/cas9 mutagenesis of SMCHD1 in HEK293Ts | /AltR1/rCrA rGrGrG rArCrC rUrUrG rCrUrU rUrUrU rArGrA rGrUrU rUrUrA rGrArG rCrUrA rUrGrC rU/AltR2/ |
| Human SMCHD1 K591A+K592A HDR donor oligo | CRISPR/cas9 mutagenesis of SMCHD1 in HEK293Ts | CATCCCATTCTATTGCTGCATATGTTGCCCAGGGACCTTGCGCTGCAGAAGGAAGATCAGGACG TGTAATTACTCCCTTAAAAAGT |
| Human SMCHD1 K668A crRNA | CRISPR/cas9 mutagenesis of SMCHD1 in HEK293Ts | /AltR1/rGrU rCrCrU rArUrC rCrArG rCrUrU rUrGrC rArArU rGrUrU rUrUrA rGrArG rCrUrA rUrGrC rU/AltR2/ |
| Human SMCHD1 K668A HDR donor oligo | CRISPR/cas9 mutagenesis of SMCHD1 in HEK293Ts | CATATTTTTTAACAGCTTTCTCAGCAACTGTCCTATCGAGCGCTGCAATTGGCACAGTTCTTACTTCATC ATATAGTGCCTGAGGT |
| Human SMCHD1 E147A F | Illumina MiSeq of clonal HEK293Ts | GTGACCTATGAACTCAGGAGTCCTGGGAGGGAAAAGTTAAGCA |
| Human SMCHD1 E147A R | Illumina MiSeq of clonal HEK293Ts | CTGAGACTTGCACATCGCAGCGCAATGGATGAAGAGATATCTGACAA |
| Human SMCHD1 K591A+K592A F | Illumina MiSeq of clonal HEK293Ts | GTGACCTATGAACTCAGGAGTCCATAAAACATTTTAAAATTCTACAGGAACAGCG |
| Human SMCHD1 K591A+K592A R | Illumina MiSeq of clonal HEK293Ts | CTGAGACTTGCACATCGCAGCGAAAAAAGTGACAATAAGTTAAACCTACCAGC |
| Human SMCHD1 K668A F | Illumina MiSeq of clonal HEK293Ts | GTGACCTATGAACTCAGGAGTCCTTTGGAAGAAACAAATGGGTGTTTG |
| Human SMCHD1 K668A R | Illumina MiSeq of clonal HEK293Ts | CTGAGACTTGCACATCGCAGCGCCCTAACAGCAGAGTAAATGAC |
