## Supplementary Table 4 for "DNA binding by ATPase-adjacent domains stimulates SMCHD1 ATPase activity"

| | Region | Disease | PMID | ATPase assay | Sex |
|---|---|---|---|---|---|
| Gln193Pro | ATPase | FSHD2 | 30698748 | dead | M |
| Leu194Phe | ATPase | FSHD2 | 31243061, 25256356 | down | M |
| His263Asp | ATPase | FSHD2 | 31243061, 25256356 | down | F |
| Tyr283Cys | ATPase | FSHD2 | 31243061, 27061275 | no effect/mild down | M |
| Tyr353Cys | ATPase | FSHD2 | 31243061, 23143600 | dead | M |
| Gly478Glu | Transducer | FSHD2 | 31243061, 25370034 | dead | M |
| Thr527Met | Transducer | FSHD2 | 31243061, 24075187 | mild down | M or F |
| Pro690Ser | Linker (loop prediction) | FSHD2 | 31243061, 23143600 | mild down | F |
